# In vivo Proximity & Spatial Proteomics with CRISPR Screening Identify STXBP1 as a Protective Modifier of α-synuclein Toxicity in Dopamine Neurons

**DOI:** 10.64898/2026.01.16.699530

**Authors:** Daichi Shonai, Julie Kent, Arinze Okafor, Yudong Gao, Pooja Parameswaran, Edwin Bustamante, Biswarathan Ramani, Yarui Diao, Martin Kampmann, Erik Soderblom, Scott Soderling

## Abstract

Parkinson’s disease (PD) is a disease of adults involving the loss of dopaminergic neurons after a long, asymptomatic, prodromal period. α-synuclein, LRRK2, and VPS35 are linked to familial PD, however, how these mutations predispose dopamine neurons to death during the early prodromal phases remains unclear. Here, we used in vivo native proximity proteomics (iBioID) and dopaminergic neuron-specific subcellular proteomics across multiple PD models to uncover early alterations preceding neuronal loss. Our analyses identified convergent disruptions in synaptic protein abundance, indicating that presynaptic trafficking defects are early events in PD pathogenesis. Using a targeted CRISPR-based genetic screen in dopamine neurons, we demonstrated that mimicking this misregulation of STXBP1 amplifies vulnerability to α-synuclein, implicating it as a previously underappreciated toxicity buffering factor. These findings highlight convergent mechanisms that sensitize dopamine neuronal degeneration and that presynaptic vesicle SNARE-complex proteins could serve as key targets for disease-modifying therapies in PD and related neurodegenerative disorders.

**Highlights:**

- In vivo native-BioID mapping of multiple Parkinson’s disease (PD) protein interactomes revealed a convergent presynaptic network.
- iBioID analysis on mutant PD proteins (α-synuclein A30P, LRRK2 G2019S, VPS35 D620N) uncovered mutation-specific shifts in local proximity networks, notably in endocytic and vesicle recycling pathways.
- Spatial proteomics (iBioCoFrac) of dopamine neurons in vivo identified functional modules with reduced levels of key synaptic proteins in PD models.
- Comparative proteomics using iBioCoFrac revealed synaptic vesicle regulation as a primary site of molecular convergence and early molecular signatures in dopamine neurons across multiple PD mouse models.
- An in vivo CRISPR screen pinpointed the presynaptic protein Stxbp1/Munc18-1 as an α-synuclein toxicity modifier in dopaminergic neurons.

## Introduction

Parkinson’s disease (PD), first described in 1817 by James Parkinson in “An Essay on the Shaking Palsy”, is the second most common neurodegenerative disorder, clinically characterized by the progressive loss of dopaminergic neurons in the substantia nigra pars compacta (SNc) and the consequent depletion of striatal dopamine^1,2^. Parkinson began his essay by acknowledging that his descriptions were based on analogy and lacked anatomical/experimental evidence. Since then, our understanding of PD has improved dramatically. Approximately 10% of PD cases follow a clear monogenic inheritance, while the vast majority (90%) are considered sporadic, underscoring intricate interactions between genetic and environmental factors^3,4^.

Notably, defects in intracellular trafficking have emerged as a central theme in PD pathogenesis^5^, with genes repeatedly implicated in vesicular transport^6–8^, lysosomal function^9,10^, mitochondrial function^11–14^, and protein quality control^15,16^ across both familial and sporadic cases. Among these, SNCA (encoding α-synuclein, α-Syn), LRRK2, and VPS35 have received particular attention^17–24^. α-Syn is tightly linked to synaptic vesicle cycling and pathological protein aggregation^25^, whereas LRRK2 and VPS35 play critical roles in lysosomal and endosomal trafficking^26–28^. Although these mutations affect distinct proteins, they converge on common pathways of vesicle trafficking and degradation, reinforcing the idea that shared molecular disruptions may underlie dopaminergic neurodegeneration^29,30^.

Despite major advances in identifying genetic risk factors and delineating their associated molecular pathways, a critical gap remains in understanding how these lifelong mutations ultimately lead to disease onset. A striking feature of PD is that individuals carrying high-penetrance mutations can remain symptom-free for decades before the eventual onset of clinical symptoms in mid-to-late adulthood, highlighting a long prodromal phase^31^. This paradox raises the possibility that subtle, early molecular alterations precede overt degeneration and establish the basis for dopaminergic neuron vulnerability. Defining whether such early alterations represent convergent molecular signatures across distinct genetic forms of PD, and whether they contribute causally to neurodegeneration, is essential for understanding disease initiation and progression.

Here, we sought to address this gap by identifying signatures of early prodromal molecular convergence underlying dopaminergic neuron vulnerability in vivo. Using in vivo HiUGE-BioID^32^ in young (6–8 weeks old) mouse models expressing wild-type or mutant α-Syn, LRRK2, and VPS35, we captured endogenous proximity proteomes for each that could drive or modulate early pathological events. These analyses revealed that all three PD-related proteins frequently converged on presynaptic factors. To evaluate whether these proximity-based hits were already altered in prodromal dopaminergic neurons, we applied dopaminergic-neuron-specific subcellular proteomics at the same early stage. By examining this predegenerative window, before overt neuronal loss or behavioral deficits, we found that several BioID-identified presynaptic proteins were misregulated in a genotype-dependent manner. These early alterations in protein localization and abundance point to molecular changes that may predispose dopaminergic neurons to later degeneration.

Through a targeted AAV-based CRISPR pooled-gRNA screen^33^ in dopaminergic neurons, we systematically tested whether convergently misregulated proteins indeed exacerbate the vulnerability of these cells to α-Syn-induced neurotoxicity. Among multiple candidates, Stxbp1 (Munc18-1) emerged as a top modifier whose loss heightened dopaminergic neuron degeneration. These results demonstrate that the proteomic alterations identified in prodromal phases can indeed exacerbate dopaminergic neuron loss, pointing to critical “conspiratorial” prodromal events in the dopamine neuronal degenerative cascade.

In summary, this multi-pronged *in vivo* approach—integrating mutant-specific proximity labeling, dopaminergic neuron-targeted proteomics, and CRISPR-based vulnerability screening provides new insight into the initial causal underpinnings of later dopamine neuronal pathogenesis that is likely relevant to PD. Our results illuminate shared trafficking disturbances in α-Syn, LRRK2, and VPS35 pathways, offering a framework that may extend to broader neurodegenerative processes involving protein aggregation, vesicular trafficking defects, and lysosomal dysfunction. Our analysis demonstrates that Stxbp1 (Munc18-1) likely functions to oppose/buffer an α-Syn toxicity suppressor gene and that its altered function in the prodromal period may ultimately contribute to neuronal loss. Modulating these early events could facilitate the future design of presymptomatic biomarkers or strategies focused at preserving dopaminergic neuron function.

## Results

### Endogenous Proximity Labeling Reveals Proteomes of Parkinson’s Risk Proteins

To map the local proximal proteomic profiles of Parkinson’s disease (PD) risk proteins in neurons *in vivo*, we employed an *in vivo* CRISPR-based BioID strategy (HiUGE-BioID) (Figure 1A and 1D)^32^. TurboID tags were integrated into each endogenous gene loci via a CRISPR-Cas9-based homology-independent universal genome engineering (HiUGE) system (Figure 1B-C)^34^, enabling the capture of both transient and stable protein interactions in proximity to the baits within the living mouse brain. This approach uniquely preserves native protein expression and regulation, thereby reducing artifacts commonly associated with conventional overexpression methods.

**Figure 1.**
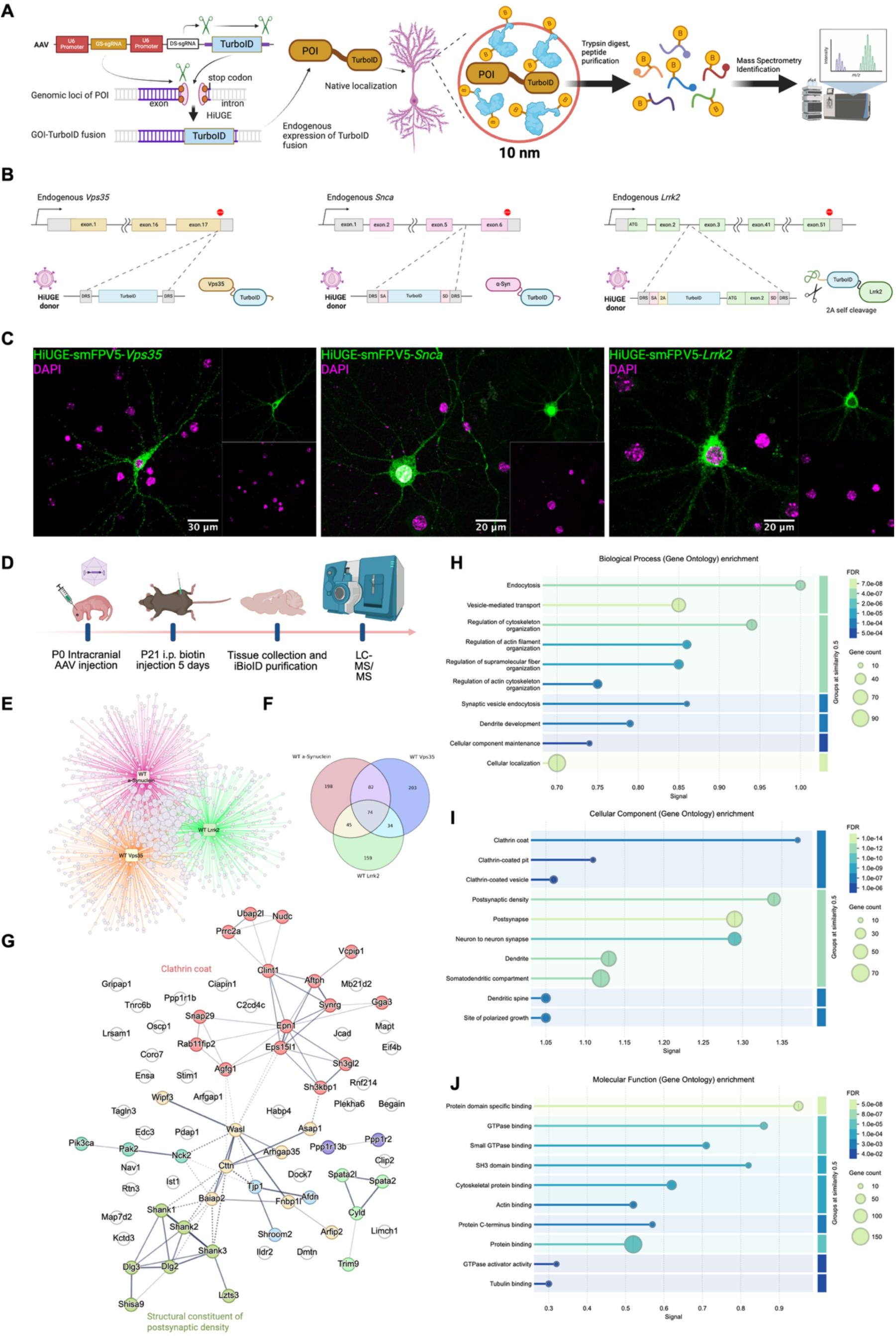
Endogenous proximity proteomics reveals molecular convergence of Parkinson’s disease risk proteins α-synuclein, Vps35, and Lrrk2 in vivo. (**A**) Schematic overview of the HiUGE-iBioID experimental workflow. TurboID was endogenously knocked into each bait gene using the HiUGE (homology-independent universal genome engineering) approach, enabling in vivo proximity labeling in native cellular environments. Following biotin administration, labeled proteins were enriched and identified by mass spectrometry. (**B**) HiUGE strategies used to insert TurboID into Snca, Vps35, and Lrrk2 loci in mice. Each schematic illustrates the TurboID donor cassette design and its integration site for each gene. (**C**) Representative immunostaining images demonstrating successful endogenous tagging of the bait proteins in cultured cortical neurons using the Spaghetti Monster v5 (smFPV5) HiUGE donor. (**D**) Overview of the in vivo experimental procedure: adeno-associated virus (AAV) injection, biotin administration, brain collection, and mass spectrometry-based proteomic analysis. (**E**) Proteomic profiles of proximity-labeled proteins for each bait (α-synuclein, Vps35, Lrrk2), highlighting the convergence of proteins identified by all three baits. (**F**) Venn diagram showing the overlap of proximity-labeled proteins detected across the three bait proteins. (**G**) Network visualization of the convergent proteins detected by all three baits, generated using STRING, with edges reflecting known or predicted protein–protein associations. (**H–J**) Gene ontology (GO) enrichment analysis of the shared proteins, showing the top 10 enriched pathways from (**H**) Biological Process, (**I**) Cellular Component, and (**J**) Molecular Function categories.

**Figure S1.**
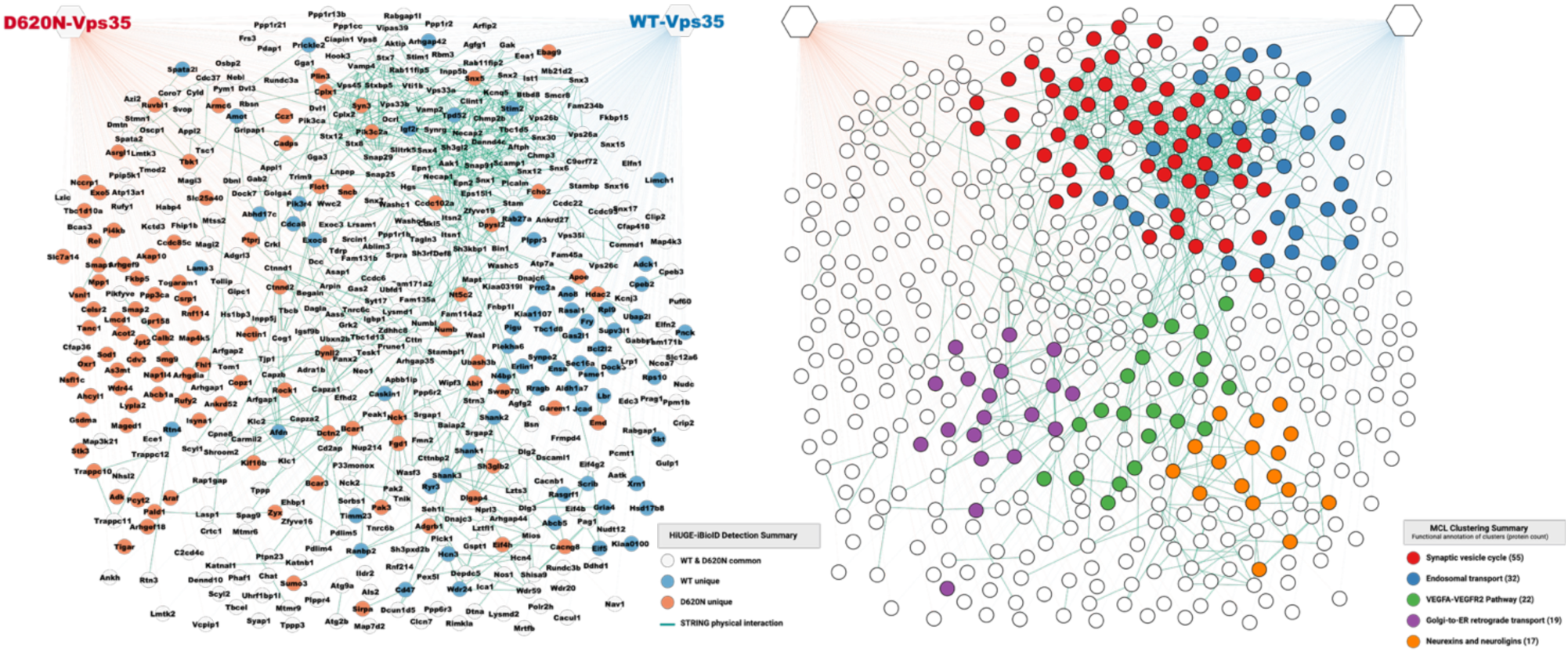
Endogenous proximity proteomics of Vps35. (**Left**) Proximity network of Vps35 in vivo from mouse brain samples in WT or D620N KI/KI mouse. Node colors represent: white, proteins identified in both conditions; blue, proteins uniquely detected in WT proximity; orange, proteins uniquely detected in D620N proximity. Edge colors indicate interaction sources: green, STRING-based physical interactions with medium confidence; blue, proximity interactions to WT Vps35 defined in this experiment; orange, proximity interactions to D620N Vps35 defined in this experiment. (**Right**) Node colors reflect functional clustering based on MCL (Markov Cluster Algorithm) with an inflation parameter of 2.0. The top five functional clusters are color-annotated.

**Figure S2.**
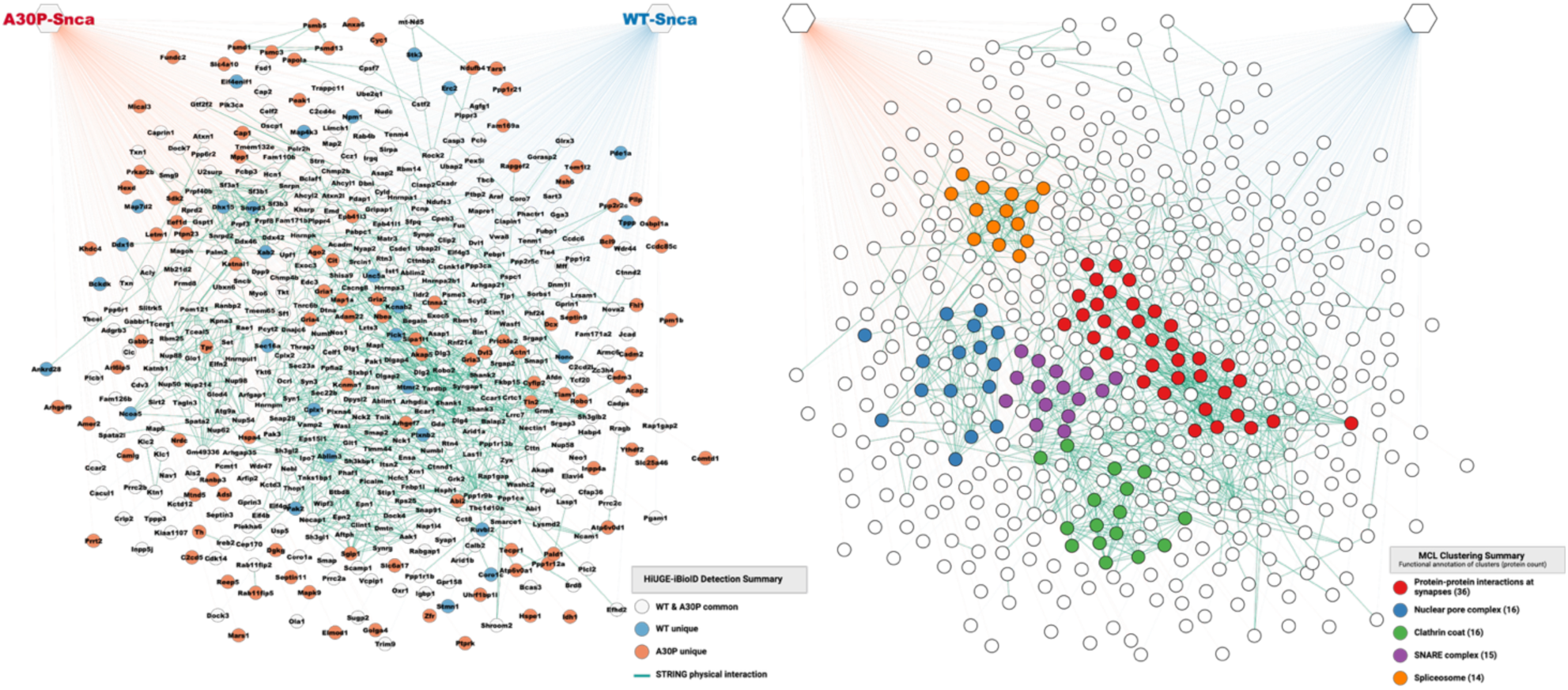
Endogenous proximity proteomics of α-synuclein. **(Left)** Proximity network of α-synuclein in vivo from mouse brain samples in WT or Α30P-Snca KI/KI mouse. Node colors represent: white, proteins identified in both conditions; blue, proteins uniquely detected in WT proximity; orange, proteins uniquely detected in A30P proximity. Edge colors indicate interaction sources: green, STRING-based physical interactions with medium confidence; blue, proximity interactions to WT α-synuclein defined in this experiment; orange, proximity interactions to A30P α-synuclein defined in this experiment. (**Right**) Node colors reflect functional clustering based on MCL (Markov Cluster Algorithm) with an inflation parameter of 2.0. The top five functional clusters are color-annotated.

**Figure S3.**
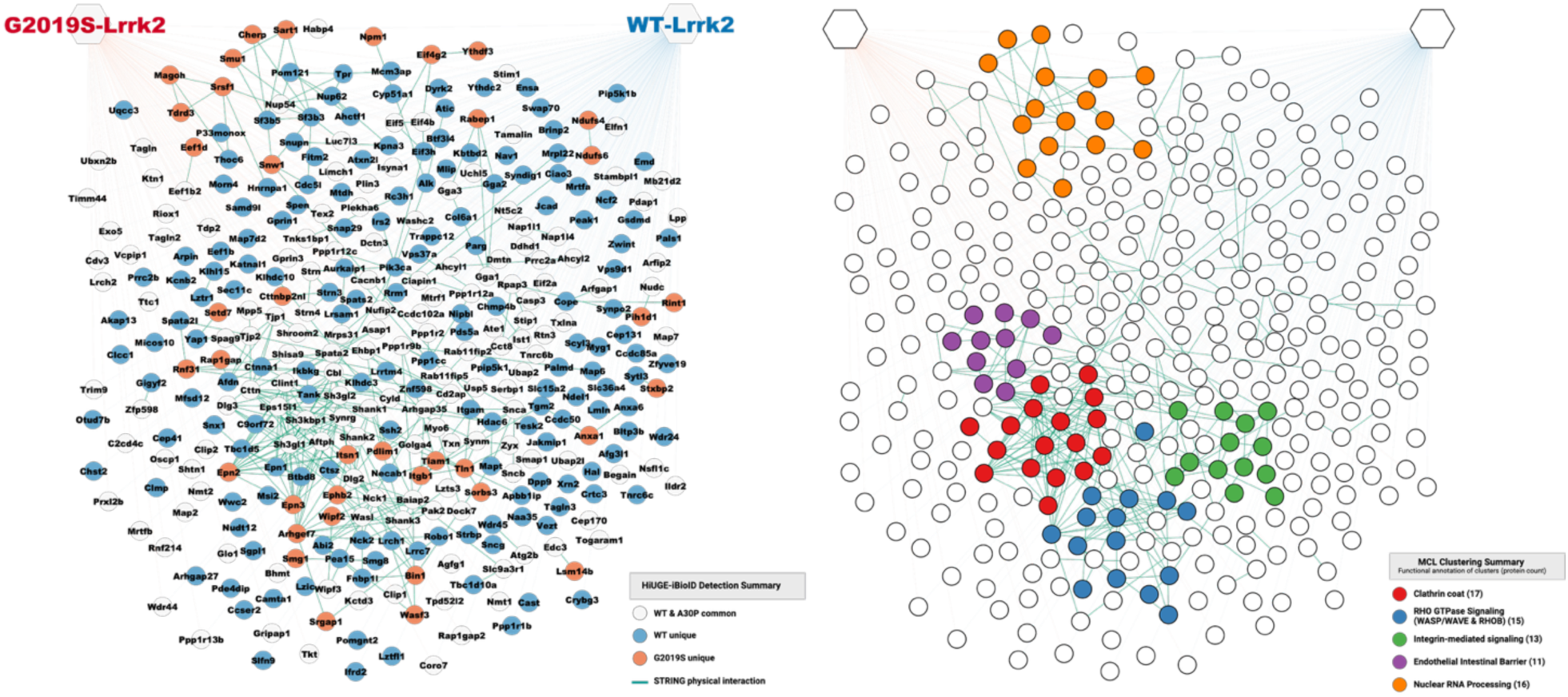
Endogenous proximity proteomics of Lrrk2. **(Left)** Proximity network of Lrrk2 in vivo from mouse brain samples in WT or G2019S-Lrrk2 KI/KI mouse. Node colors represent: white, proteins identified in both conditions; blue, proteins uniquely detected in WT proximity; orange, proteins uniquely detected in G2019S proximity. Edge colors indicate interaction sources: green, STRING-based physical interactions with medium confidence; blue, proximity interactions to WT Lrrk2 defined in this experiment; orange, proximity interactions to G0219 Lrrk2 defined in this experiment. (**Right**) Node colors reflect functional clustering based on MCL (Markov Cluster Algorithm) with an inflation parameter of 2.0. The top five functional clusters are color-annotated.

The validity of our approach was assessed through LC-MS/MS analyses of the wild-type (WT) proximity proteomes for known interactors. We employed a PHP.eB-packaged HiUGE adeno-associated virus to integrate TurboID into 4-week-old Cas9-expressing mice, with an IRES-TurboID HiUGE donor serving as a negative control that expressed soluble, non-fusion TurboID. Following biotin administration, we purified the biotinylated proteins and performed LC-MS/MS analyses. Data were normalized against the soluble TurboID controls, and significantly enriched proteins were defined as those with log₂ fold change > 1 and p < 0.05 (two-tailed heteroscedastic t-test; three biological replicates).

For Vps35, we identified 322 significantly enriched proteins, including core retromer components (Vps26a, Vps26b) sorting nexins (Snx1, Snx2, Snx3, Snx4), and key subunits of the WASH complex (Washc1, Washc4, Washc5), consistent with its known function in endosomal recycling (Figure S1) ^35–37^. Similarly, α-synuclein was enriched for known presynaptic interactors such as Synapsin 1, Synaptobrevin-2 (VAMP2), and Endophilin A1, as well as RNA-binding proteins like Sfpq, Matrin-3, Nucleophosmin and Rbm14, reflecting its roles in synaptic and nuclear functions^38–41^ (Figure S2). The Lrrk2 proximity proteome comprised 311 enriched proteins, notably featuring proteins associated with vesicular transport and cytoskeletal regulation, including retrograde transport regulators (Snx1, Tbc1d5)^42^, RAB GTPase interacting proteins (Rab11fip2, Rab11fip5) and Auxilin (Dnajc6), a known PD-associated protein phosphorylated by Lrrk2 (Figure S3)^43^.

Collectively, these findings confirm the physiological relevance of our endogenous TurboID proximity labeling approach to define proximity proteomic maps for α-synuclein, Vps35, and Lrrk2. This dataset established the foundation for subsequent analyses focused on convergent molecular interactions and genotype-specific proximity alterations relevant to Parkinson’s disease pathogenesis.

### Convergent Molecular Interactions of PD-Related Proteins with Synaptic Processes

To systematically delineate the shared molecular proximity networks among wild-type α-Synuclein, Lrrk2, and Vps35, we performed a comparative analysis of the proximity proteomic data obtained from the HiUGE iBioID experiments. Our analysis identified a “convergent proteome” consisting of 74 proteins consistently detected across all three baits (three-way hypergeometric test, p = 0.0026), along with an additional 161 proteins observed in at least two datasets (Figure 1E-F). This robust overlap underscores significant molecular convergence among these PD-associated proteins.

This convergent proteome prominently features regulators of presynaptic vesicle dynamics, many of which also contribute to endocytosis and vesicle trafficking. Key proteins include Rab11 Family Interacting Protein 2 (Rab11fip2), Asap1, Endophilin-A1 (Sh3gl2), Wasl (Wiskott-Aldrich Syndrome Protein-like), and WAS/WASL-interacting protein family member 3 (Wipf3) (Figure 1G). Notably, proteins specifically linked to synaptic vesicle recycling were also found, such as Eps15L1, Epsin-1 (Epn1), and Clathrin interactor 1 (Clint1). The convergent proteome was also enriched in synaptic components. Presynaptic proteins involved in vesicle docking and trafficking were represented, including Snap29, Pacsin1 (Syndapin-1), and Trim9 (Tripartite motif-containing 9), reinforcing the presynapse as a key site of convergence among these PD risk proteins. Several postsynaptic proteins, including Shank1, Shank2, Shank3, and Syngap1, were also enriched. Gene Ontology analysis further confirmed significant enrichment for biological processes related to vesicle-mediated transport and synaptic vesicle cycles, as well as the cellular component of clathrin-coated vesicles (Figure 1H-I). Furthermore, Gene Ontology molecular function analysis revealed significant enrichment for proteins involved in GTPase regulation, providing functional insight into the convergence of these PD-related proteins within synaptic contexts (Figure 1J). Altogether, these data show that α-Synuclein, Lrrk2, and Vps35 converge on clathrin-mediated endocytosis particularly in processes governing synaptic vesicle turnover and trafficking, indicating a mechanistic link between these processes and PD-associated synaptic dysfunction.

### Mutant-Specific Proximity Proteomes of Parkinson’s Disease Proteins

Previous studies have shown that individual missense mutations in PD-risk proteins alter their molecular features^44–54^. However, the effect of such single amino acid substitutions under native in vivo brain conditions has not been systematically examined. Here, to elucidate genotype-specific alterations in proximity proteomes associated with Parkinson’s disease (PD) mutations in its native state, we generated double homozygous knock-in models (*KI-Vps35^D620N/D620N^, KI-Snca^A30P/A30P^, KI-Lrrk2^G2019S/G2019S^*)^55,56^ crossed with a line of mice constitutively expressing *Streptococcus pyogenes* Cas9 (*H11-Cas9* line) (Figure 2A). Using the same strategy applied for WT proximity proteomes, we performed LC-MS/MS analyses normalized against soluble TurboID controls. Significantly enriched proteins were defined as those with log₂ fold change > 1 and p < 0.05 (two-tailed heteroscedastic t-test; three biological replicates). This analysis identified 357, 473, and 244 proximity proteins for D620N Vps35, A30P α-synuclein, and G2019S Lrrk2 mutants, respectively (Figures S1-S3). Notably, substantial overlaps with wild-type (WT) proteomes were observed for Vps35 (71.9%), and α-synuclein (74.6%), and less so for Lrrk2 (35.7%). Overall, there was high concordance between genotypes in Gene Ontology (GO) terms (85.05% biological processes, 79.59% cellular components, 76.47% molecular functions) (Figure S4). This overlap highlights core proximity networks despite disease-associated mutations, while also uncovering selective, mutation-driven neomorphic associations.

**Figure 2.**
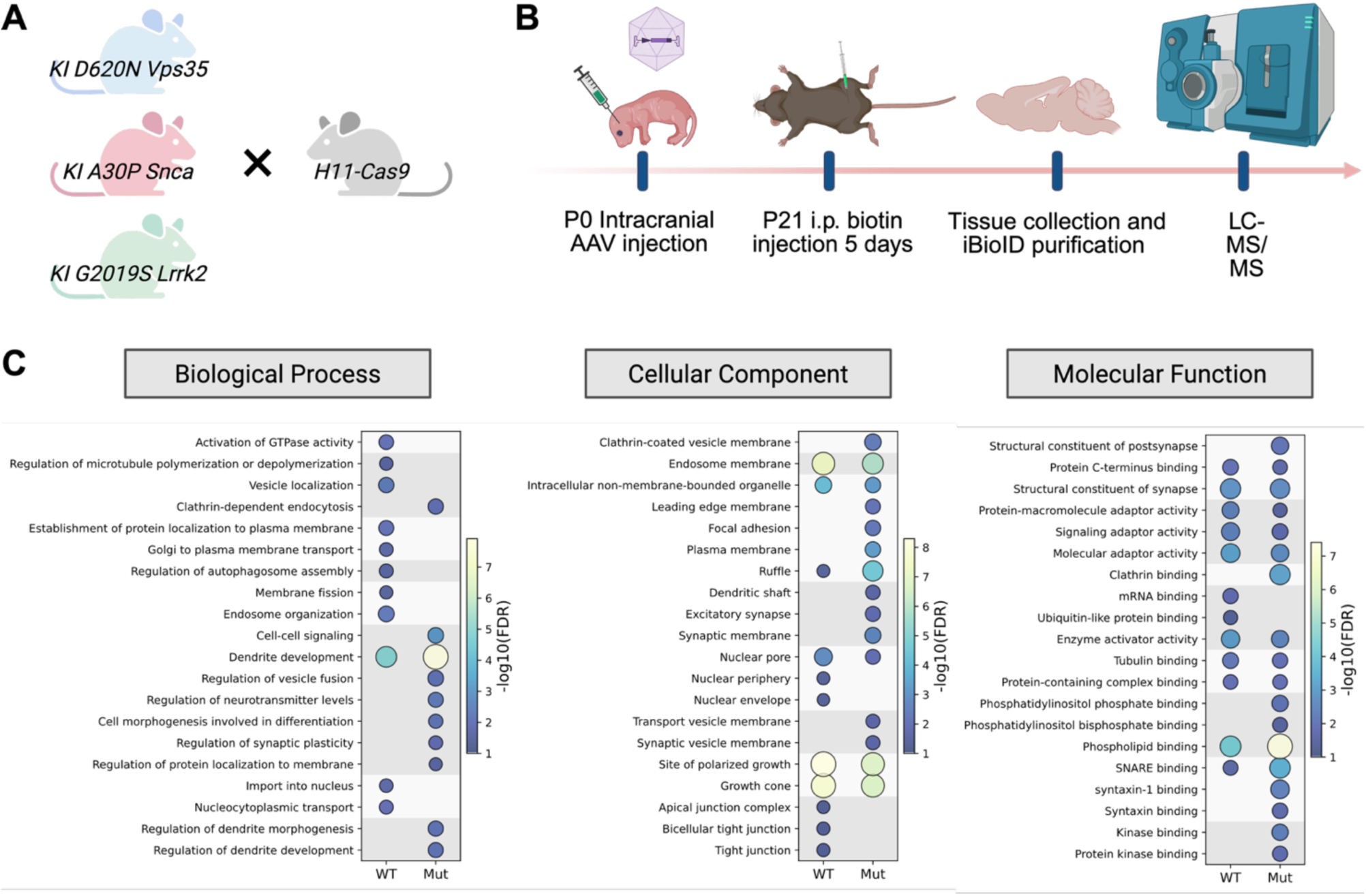
Genotype-Specific Proximity Proteomes Reveal Mutation-Driven Alterations in PD Risk Mutants. **(A)** Mouse strain strategy for mutant-specific HiUGE-iBioID. For in vivo analysis of three PD-related proteins—D620N Vps35, A30P Snca, and G2019S Lrrk2—each mutant line was crossed to generate homozygous PD-mutant KI/KI; Cas9+/- mice, which were used for the HiUGE-iBioID experiment. **(B)** Timeline of the HiUGE-iBioID experiment. **(C)** Comparison of GO analysis results (Biological Process, Cellular Component, Molecular Function) between the convergent WT proximity proteome and the convergent mutant proximity proteome. Gene lists were generated based on proteins detected in at least two out of three bait proteins, separately for WT and mutant. The top 10 and bottom 10 GO terms showing the largest differences were selected based on enrichment signal (FDR and fold change). Each ontology was subjected to hierarchical clustering based on Jaccard similarity of the component genes and is separated in the plot accordingly. The color of each node reflects the −log10(FDR), and the marker size represents the enrichment signal.

**Figure S4.**
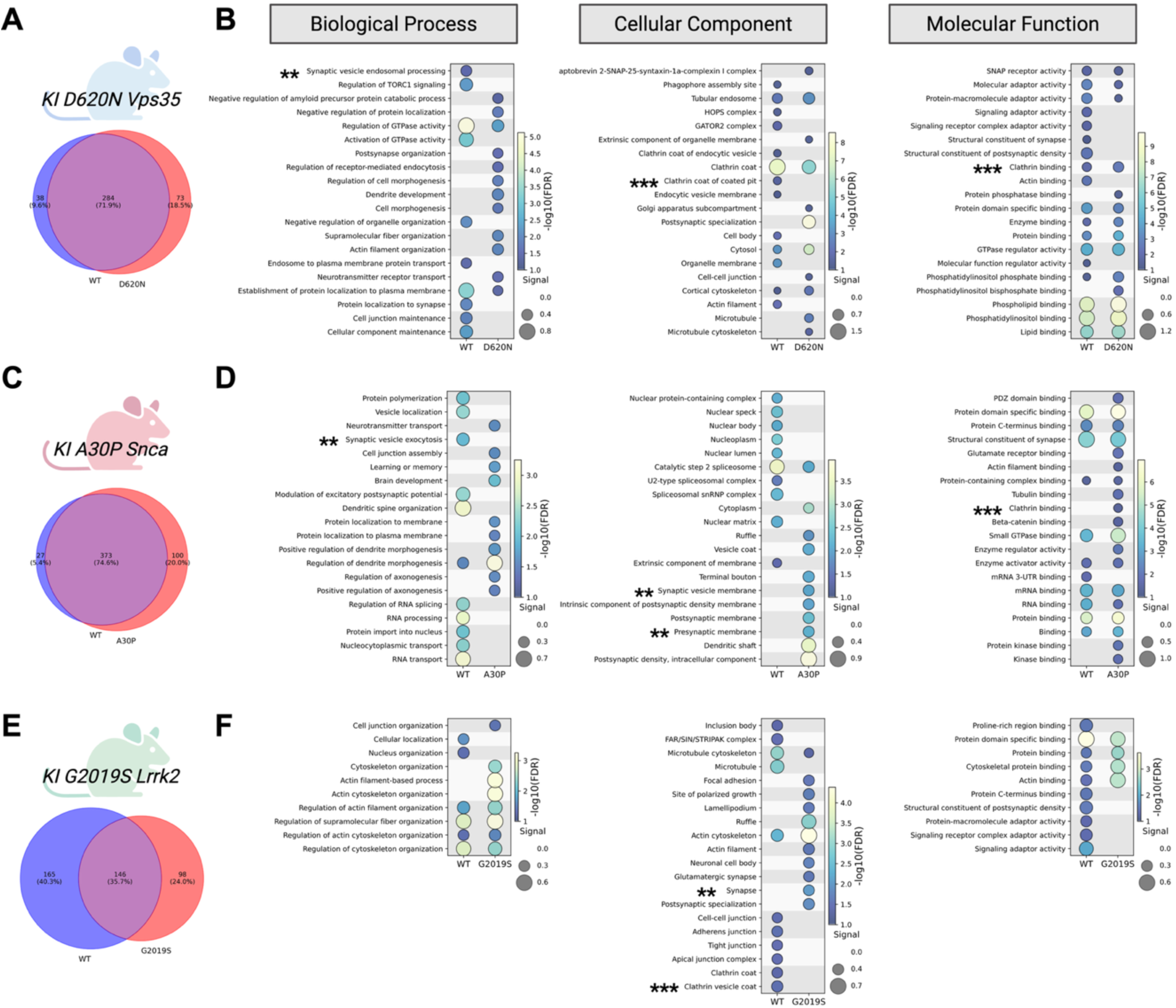
Comparative analysis of the proximity proteomes of WT and mutant forms of Parkinson’s disease–associated proteins. **(A, C, E)** Venn diagrams showing the overlap of proximity-labeled proteins detected in WT and PD mutant forms of Vps35, Snca, and Lrrk2, respectively. **(B, D, F**) Comparative gene ontology (GO) analysis for each corresponding bait. For Vps35 and Snca, the top 10 and bottom 10 GO terms showing the largest differences between WT and mutant proximity proteomes were selected based on enrichment signal (FDR and fold change). For the Lrrk2 experiment, the top 5 and bottom 5 GO terms were selected. Each ontology was subjected to hierarchical clustering based on Jaccard similarity of the component genes and is separated in the plot accordingly. The color of each node reflects the −log10(FDR), and the marker size represents the enrichment signal.

### Convergent Disruptions in Clathrin-Mediated Endocytosis and Vesicle Recycling

A focused investigation of the mutant proteomes uncovered a pronounced enrichment of proteins involved in clathrin-mediated endocytosis and vesicle recycling, pathways previously linked to PD ^57,58^ (Figure 2C). The D620N Vps35 mutant specifically enriched endocytic regulators (e.g., Numb, Endophilin-B2 (SH3GLB2), Smap1, Smap2, Fcho2), suggesting altered dynamics in clathrin-coated pit formation (Figures S1 and S4). The A30P α-synuclein mutant displayed enhanced proximity to regulators of endosomal trafficking such as Rab11fip5, Acap2, Sgip1, and Annexin A6, implicating altered recycling endosome pathways (Figures S2 and S4). G2019S Lrrk2 mutant show neomorphic proximity association with Annexin A1, Epsin 2, Epsin 3, as well as two effector proteins of Rab4/5, Rabenosyn-5 (Rbsn) and Rabaptin-5 (Rabep1)(Figures S3 and S4). Together, these mutation-specific alterations in the in vivo brain proximity networks implicate a shared alteration of intracellular trafficking machinery across PD-linked mutations. Previous studies have also suggested defects in clathrin-mediated endocytosis and vesicle recycling related to Parkinson’s disease associated phenotypes^8,57–62^. Our findings provide an unbiased, proteome-wide validation that PD-associated mutations are likely to systematically rewire endocytic and vesicle recycling networks.

### Alterations in Synaptic Vesicle Trafficking and SNARE Complex Interactions

Comparative analyses revealed specific shifts in proximal proteins that affect synaptic vesicle trafficking and SNARE complex dynamics (Figure 2C). The A30P α-synuclein mutant exhibited altered proximity to SNARE-related proteins, notably forming neomorphic proximity associations with Prrt2 (Proline-rich transmembrane protein 2), while losing association with Complexin-1 (Cplx1), a key mediator of vesicle priming and fusion, which is detected in WT α-synuclein proximity. Together this suggests altered vesicle fusion specificity. Unique detection of Atp6v0a1^63^ and Atp6v0d1 in proximity to A30P α-synuclein implies potential changes in vesicular acidification and neurotransmitter loading. Furthermore, this mutant showed preferential associations with presynaptic vesicle regulators, indicating a reorganization of the presynaptic protein network. In parallel, the D620N Vps35 exhibited a neomorphic interaction with NSFL1 cofactor p47 (Nsfl1c), a key regulator of SNARE disassembly. It also showed increased proximity to Cplx1, β-synuclein, and Synapsin-3, while losing associations with Rab27a and Exoc8, which coordinate synaptic vesicle tethering and fusion. Lastly, the G2019S Lrrk2 mutant demonstrated preferential proximity to Stxbp2 (Munc18-2) and Synaptotagmin-4, reinforcing its broad impact on SNARE-mediated vesicle docking and neurotransmitter release (Figure S1 – S4). These observations align with prior reports that LRRK2 mutations affect dopamine release and synaptic dynamics^64–66^, thereby contributing to PD pathogenesis.

### Development and Validation of iBioCoFrac for in vivo Dopaminergic Neuron Unbiased Co-Fractionation Proteomics

Using the HiUGE-iBioID approach, we first demonstrated that synaptic vesicles and clathrin-dependent endocytosis act as convergent points for PD-associated proteins, suggesting local vulnerability to missense mutations. These observations raised a critical systems-level question: beyond local disruption, how do PD mutations rewire the subcellular architecture of dopamine neurons? PD-related proteins such as Vps35, α-synuclein, and LRRK2 participate in numerous intracellular trafficking pathways, notably those associated with the endo-lysosomal system. Furthermore, PD is linked to dysfunction in various organelles, including mitochondria, autophagic vesicles, and granule bodies, highlighting the complexity of the disease’s cellular impacts^6–16^. Importantly, it remains unclear whether these organelle dysfunctions represent primary, convergent mechanisms or secondary outcomes of broader cellular disturbances.

To systematically map these alterations, an unbiased, cell type–specific spatial proteomics method was required. Existing co-fractionation proteomics approaches^67^ leverage gentle lysis of cultured cells and subcellular fractionation with quantitative mass spectrometry to simultaneously analyze multiple compartments within complex biological samples. These methods infer subcellular localization by identifying proteins whose abundance patterns co-vary across fractions, enabling the detection of spatially organized protein modules. Our previous study^68^ extended this strategy to the tissue, demonstrating that genetic mutations induce large-scale reorganization of neuronal subcellular proteomes. However, a critical limitation remained: these methods lacked the ability to resolve changes within genetically defined cell types.

To overcome this limitation, we developed iBioCoFrac, an in vivo cell type-specific biotin-enriched co-fractionation proteomics strategy (Figure 3A-I). This innovative method integrates cell type-specific, unbiased proteome biotin labeling with systems-level spatial proteomics, profiling co-fractionated protein networks specifically within dopamine neurons. To our knowledge, this represents the first application of a co-fractionation spatial proteomics workflow with cell type specificity in vivo, providing physiologically relevant spatial proteomic insights into neuronal contexts. iBioCoFrac, therefore, provides a framework for mapping subcellular proteomes in genetically defined neuronal populations in vivo.

**Figure 3.**
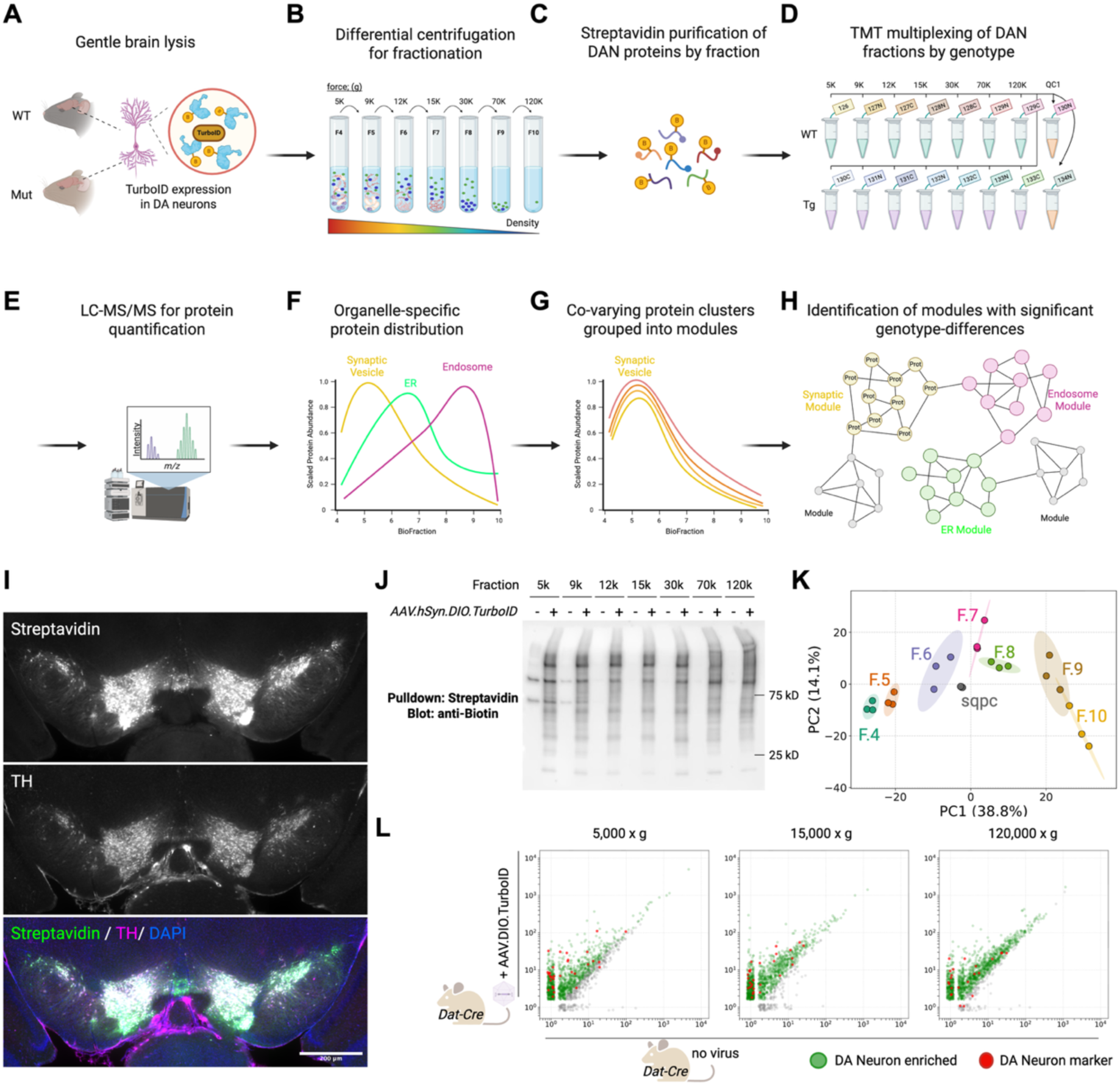
iBioCoFrac Workflow Enables Dopamine Neuron-Specific Unbiased Subcellular Spatial Proteomics. **(A–L)** Schematic overview of the iBioCoFrac workflow for selective unbiased subcellular proteomic profiling of dopamine neurons in vivo. **(A)** To map the dopamine neuron proteome, soluble TurboID was expressed in a dopamine neuron–specific manner using the Dat-Cre mouse line. **(B)** Brain tissues underwent differential centrifugation for subcellular fractionation. **(C)** Streptavidin enrichment was performed for each fraction to isolate biotinylated proteins. **(D)** TMT multiplexing was applied to the streptavidin-purified samples to compare protein abundance across genotypes. **(E)** Proteins were quantified by LC-MS/MS. **(F)** Protein abundance patterns across the fractions were used to build a covariance matrix. **(G)** Proteins with similar distribution patterns were grouped into co-varying modules using Pearson correlation and the Leiden algorithm. **(H)** Each detected module was annotated based on gene enrichment analysis and used to identify modules with significant genotype-dependent differences. **(I**) Immunofluorescence images demonstrating dopamine neuron–specific biotinylation, visualized by streptavidin staining. **(J)** Western blot analysis with anti-biotin antibody showing enrichment of biotinylated proteins from dopamine neuron fractions after co-fractionation. **(K**) Principal component analysis (PCA) plot illustrating distinct proteomic profiles of the fractionated samples. **(L)** Mass spectrometry reveals significant enrichment of dopamine neuron proteins ^69^ in the isolated fractions.

In our iBioCoFrac strategy, cell type specificity is achieved by expressing the promiscuous biotin ligase TurboID exclusively in dopamine neurons using an AAV-hSyn-DIO-TurboID construct delivered to Dat-Cre mice (Figure 3A). Building on our previously established in vivo workflow^68^, we performed subcellular fractionation (Figure 3B), followed by streptavidin-based enrichment of biotinylated dopaminergic proteins (Figure 3C). The enriched proteins were then digested, labeled with 16-plex TMT reagents, and quantified by LC-MS/MS. This effectively allowed for the quantification of dopamine neuronal protein abundance across genotypes from discrete subcellular fractions (Figure 3D and 3E). In doing so, it also reveals co-fractionated protein networks that are physically and functionally associated (Figure 3F-H). We thus reasoned that iBioCoFrac could bridge the gap between local protein interactions detected by proximity labeling and broader cellular-scale dysfunctions, elucidating how missense mutations disrupt both local and network-level protein organization.

To validate the iBioCoFrac workflow, we first confirmed its ability to accurately quantify protein abundance in dopamine neurons–enriched samples. E15.5 mice underwent in utero intracranial AAV injections, followed by daily biotin administration for five consecutive days at two months of age (Figure 3I and Figure 4A). Brain regions, including the substantia nigra pars compacta and striatum, were dissected and fractionated. Biotinylated proteins from each fraction were enriched via streptavidin purification (Figure 3J), TMT labeled, and analyzed by LC-MS/MS. PCA plots of these validation experiments demonstrated distinct patterns of biotinylation for each fraction and enrichment of known dopamine neuron–derived proteins, such as Sodium-dependent dopamine transporter (DAT, Slc6a3), Tyrosine hydroxylase (Th) and Aromatic-L-amino-acid decarboxylase (Ddc), across biological replicates (Figure 3K and 3L)^69^. Based on the technical reliability and fidelity of iBioCoFrac for mapping neuronal subcellular proteomes, this approach was next applied to α-syn, Vps35, and Lrrk2 wildtype and mutant mouse models.

### iBioCoFrac Reveals Convergent Alterations in Synaptic Vesicle and Clathrin-Associated Proteins

For investigating the subcellular proteomic landscape with our newly developed iBioCoFrac pipeline, we employed three mouse models containing PD-relevant mutations—*KI D620N Vps35*^56^, *Tg-Th-hSnca^A30P/A53T^* ^70^, and *Tg-Pdgfb-hLRRK2^G2019S^* ^71^ (Figure 4A). Although each model exhibits only late-onset or subtle neurodegenerative phenotypes (with limitations such as mild neuron loss or variable pathology)^72–76^ their combined use has the potential to investigate early molecular changes downstream of genetic risk factors. By crossing these mouse lines with *Dat-Cre* mice, we enriched for dopamine neurons in our fractionated iBioCoFrac samples. We reasoned this intersectional strategy could leverage the complementary strengths of each model to reveal common mechanisms underlying early pathogenic alterations in Parkinson’s disease.

In our initial gene-level analysis, we applied a protein-by-protein two-way repeated measures ANOVA to assess genotype effects across subcellular fractions (Figure S5). Several PD-linked proteins were altered in each of three models. Snca and Dnajc6 were downregulated in the *KI D620N Vps35*; Vps35, Synj1, and Eif4g1 were downregulated in the *Tg-Pdgfb-hLRRK2^G2019S^*; and Eif4g1 was also affected in the *Tg-Th-hSnca^A30P/A53T^* model. This approach also revealed significant alterations in proteins associated with synaptic vesicles and clathrin-coated membranes across all three PD models. Notably, common trends emerged despite some model-specific nuances. For example, in the *Tg-Pdgfb-hLRRK2^G2019S^*^S^ model, Rab3c, a known substrate of LRRK2^77^, displayed a significant upregulation, whereas Rab8b was selectively upregulated in the *KI D620N Vps35* model. Additionally, the clathrin regulator Cltc was misregulated in both the *Tg-Pdgfb-hLRRK2^G2019S^*and *Tg-Th-hSnca^A30P/A53T^* models, and Hspa8, implicated in clathrin uncoating and protein misfolding, was consistently downregulated in the *Tg-Pdgfb-hLRRK2^G2019S^* and *KI D620N Vps35* models, highlighting clathrin-related regulation as a key affected process. Several v-ATPase subunits (Atp6v1c1, Atp6v1e1, Atp6v1h) and neuronal SNARE-related proteins (Stxb1, Syp, Nsf) also exhibited altered abundance, underscoring a potential effect of genotype on vesicle dynamics. Together, these findings demonstrate that proteins enriched by proximity proteomics of PD risk factors also exhibit altered abundance in vivo within dopamine neurons across multiple genetic models of Parkinson’s disease.

**Figure S5.**
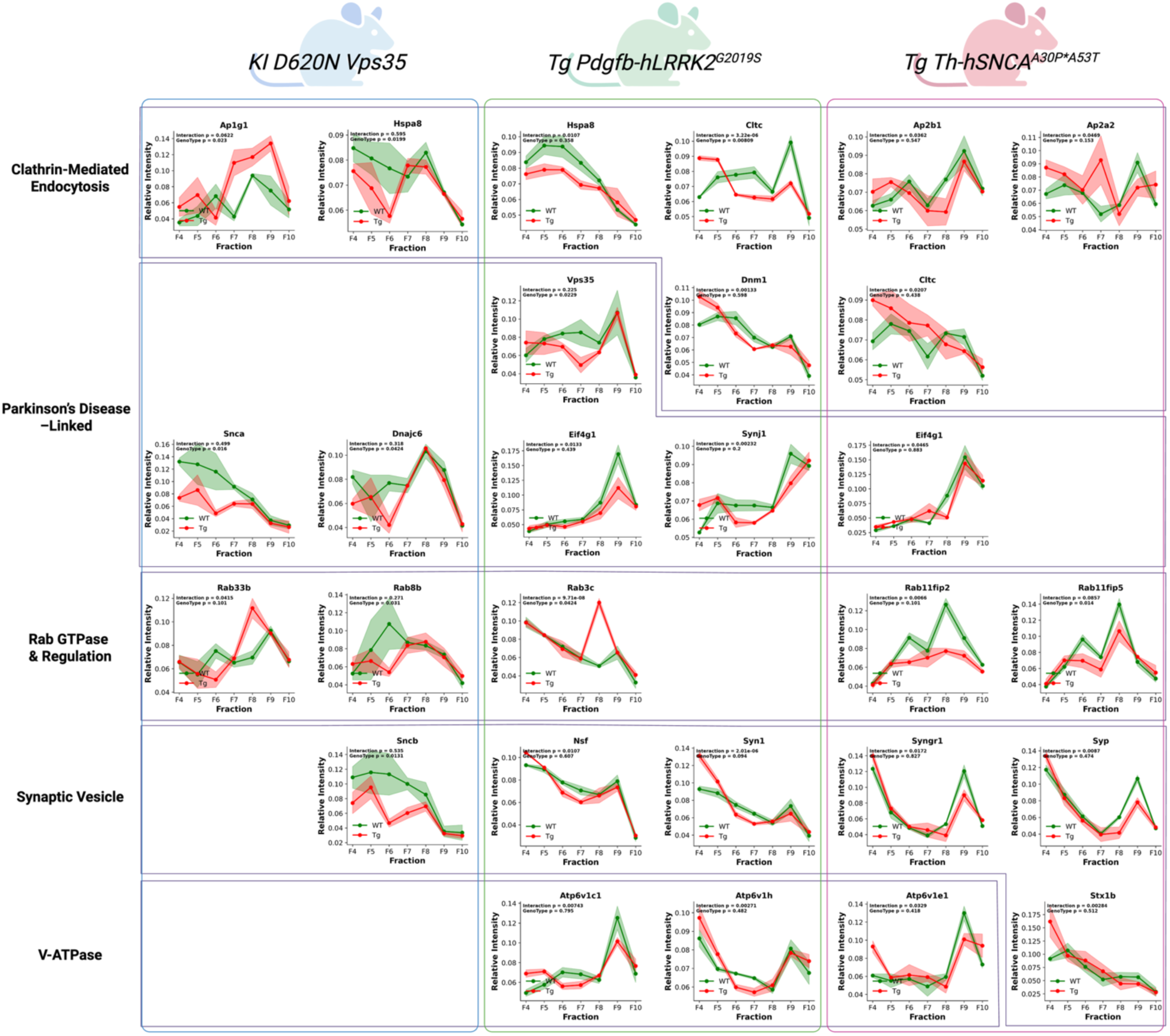
Protein abundance changes in dopamine neurons across genetic mouse models of Parkinson’s disease. Protein abundance was quantified across iBioCoFrac fractions, and genotype-dependent effects were evaluated using a protein-by-protein two-way repeated measures ANOVA. Proteins showing either significant genotype × fraction interactions or significant genotype main effects are displayed. Proteins are organized by the genetic mouse model in which the alteration was detected, and their associated biological processes.

### Covariance Analysis of Dopamine Neuron-Enriched Proteomes in PD Models

We next extended our analysis beyond individual protein abundance to a network-level perspective to gain more biologically meaningful insights into how PD genetic risk factors perturb the neuronal subcellular proteome. Using covariance analysis, we grouped proteins that co-vary across subcellular fractions into functional modules representing predicted subcellular compartments. This bottom-up, data-driven approach allowed us to evaluate systematic differences between wild-type and mutant dopamine neuron-enriched proteomes at the level of entire protein modules and pathways rather than individual proteins. For clustering, we employed the Leiden algorithm ^78^, which identifies connected protein modules and converges to high-quality, locally optimal partitions. the goal of this network-level analysis was to place differentially abundant proteins into a meaningful biological context, complementing the protein-by-protein analysis.

Using network analysis of the iBioCoFrac data, we identified covarying protein modules across three distinct PD models. This analysis revealed 19, 16, and 39 unique modules in the *KI D620N Vps35*, *Tg-Th-hSNCA^A30PA53T^,* and *Tg-Pdgfb-hLRRK2^G2019S^*models, respectively (Figure 4B-4D). These modules, represent clusters of proteins with highly correlated co-fractionation profiles, suggesting each module contains shared subcellular and/or functional interactions. Subsequent statistical analysis using an extended MSstatsTMT linear mixed-model framework ^79^(with Bonferroni correction (p-adjust < 0.05) identified 12 significantly altered modules in the *KI D620N Vps35* model, 7 in the *Tg-Th-hSNCA^A30PA53T^*model, and 14 in the *Tg-Pdgfb-hLRRK2^G2019S^* model under transgenic versus wild-type conditions. Among these significantly altered modules, we identified several enriched for proteins involved in synaptic vesicle regulation: Module 7 in *KI D620N Vps35*, Module 4 in *Tg-Th-hSNCA^A30PA53T^*, and Module 11 in *Tg-Pdgfb-hLRRK2^G2019S^*, maintaining a consistent pattern across models (Figure 5A) These modules were commonly downregulated at a significant level across all models, with the most pronounced effect observed in Module 11 in *Tg-Pdgfb-hLRRK2^G2019S^* (Figure 5D). Like the proximity proteomics analysis, several proteins in these modules, such as Snca, Snap91, Bin1, and Hspa8, regulate synaptic vesicle functions (Figure 5B and E). Additionally, Snca, Sncb, and Mapt, are implicated in neurodegeneration^80,81^ and are also associated with synaptic dysfunction, while Slc6a3, a dopamine transporter, modulates dopamine neurotransmission. Their presence within these modules reinforces their contribution to synaptic vesicle dysregulation observed in these PD models. Notably, these alterations were detected as early as two months of age, emphasizing early network-level disruptions that precede overt neurodegeneration.

**Figure 4.**
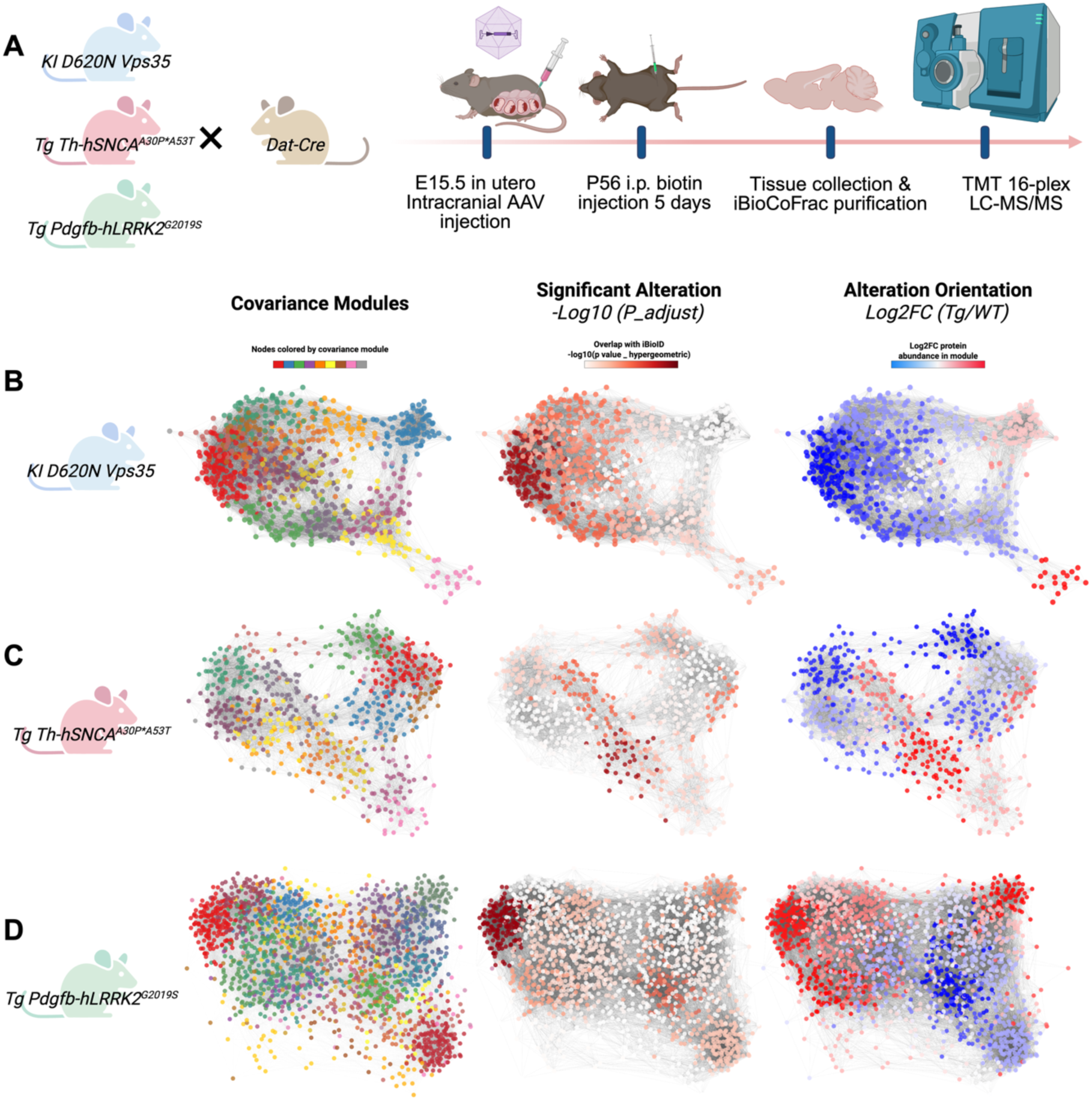
Comparative Subcellular Spatial Proteomics of Dopamine Neurons in Parkinson’s Disease Models Using iBioCoFrac. (**A**) Schematic overview of the iBioCoFrac workflow for comparing subcellular spatial proteomes of dopamine neurons between Parkinson’s disease (PD) models and wild-type controls. For each group, tissue from five animals was pooled per biological replicate (n=3 per group). (**B-D)** Module detection results for *KI D620N Vps35*, *Tg-Th-hSnca^A30P/A53T^*, and *Tg-Pdgfb-hLRRK2^G2019S^*Parkinson’s disease missense mutant models, respectively. Modules were identified using the Leiden algorithm with network enhancement. The statistical significance of module-level alterations was determined using Bonferroni correction, and module-level changes are depicted as log₂ fold changes between PD model and wild-type samples. Networks were visualized in Cytoscape (v3.10.3) using a Prefuse Force Directed layout, where nodes represent proteins and edges indicate significant covariation (Network Enhancement score > 1.0) between protein abundance profiles. Edge weights were proportional to NE scores, and weak edges below the threshold were excluded for clarity. Node colors correspond to module assignments (left), statistical significance (−log₁₀ P_adjust, middle), or fold-change direction (log₂FC, right).

### Module enriched for proximity proteome components reveals synaptic proteins as early molecular signatures of dopamine neuronal dysfunction

We next analyzed the dysregulated modules of each genotype for an enrichment of proteins within proximity to Vps35, α-synuclein, and LRRK2. We reasoned these modules would represent higher confidence modulators of dopamine neuronal dysfunction for functional analysis. To pinpoint these candidate protein modules, we first performed an unbiased comparison between the protein modules derived from iBioCoFrac and the proximity networks of mutated Vps35, α-synuclein, and LRRK2 using a hypergeometric test (Figure 5A). Among all the modules, module M11 emerged as particularly significant (Figure 5B). This module, comprising 77 proteins, showed a striking overlap with the PD proximity networks: 42.9% of its proteins were associated with the Vps35 network, 53.2% with the α-synuclein network, and 29.9% with the LRRK2 network (individual proteins may belong to more than one network) (Figure 5C).

**Figure 5.**
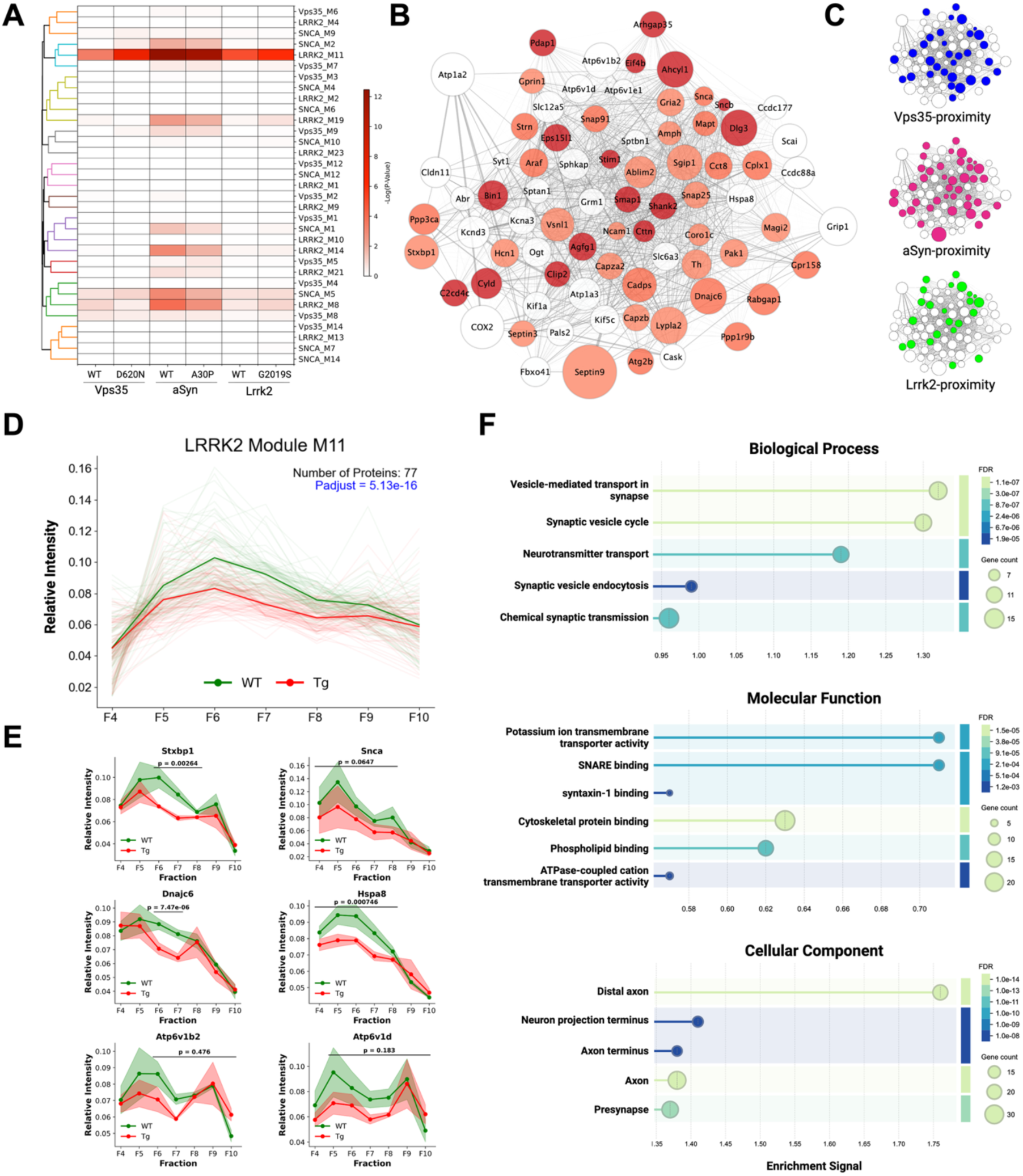
Comparative Proteomics Identifies Synaptic Compartments as Early Molecular Signatures of Parkinson’s Disease. (**A**) Comparison of iBioCoFrac-defined biological modules with iBioID proximity proteomics results performed independently in this study. Each cell is color-coded to reflect the hypergeometric p-value indicating the extent of overlap between all modules and the entire iBioID dataset. (**B**) Network visualization of Module 11 identified in the Lrrk2^G2019S^ transgenic model. Node color reflects the frequency of proximity detection by the three PD protein baits (Vps35, α-synuclein, and Lrrk2) in the iBioID experiment (1, 2, or 3 times). Edges represent STRING (Mus musculus) network interactions at medium confidence (0.40). All 77 Module 11 proteins are shown. (C) Separate network visualizations showing proximity proteins in Module 11 detected by each of Vps35, α-synuclein, or Lrrk2 baits in the iBioID experiment. (D) Comparison of protein abundance in Module 11 (WT-MUT Contrast log2Fold-Change=−0.18, T=−8.60, DF=3144, p-adjust=5.13×10−16; n=3 independent experiments). (E) Line plots showing individual protein abundance across fractions (F4 to F10) for all Module 11 proteins. (F) Gene ontology (GO) enrichment analysis of Module 11 proteins, showing results for Biological Process, Cellular Component, and Molecular Function domains, as determined by STRING API with FDR < 0.05.

Further analysis revealed that M11 is significantly downregulated in dopamine neurons in the Tg-Pdgfb-hLRRK2 model (Log₂FC = –0.18, p = 1.04E–18), suggesting that PD risk factors may mediate local alterations in protein proximity networks that lead to specific changes in protein abundance (Figure 5D). Gene ontology enrichment analysis of M11 highlighted overrepresentation of synaptic functions, with significant enrichment in terms such as “synaptic vesicle cycle” (GO:0099504), “distal axon” (GO:0097458), and “presynaptic active zone” (GO:0048786) (Figure 5F). In addition, M11 includes several proteins previously implicated in PD, such as α/β -synuclein, tau (Mapt), and auxilin (Dnajc6), as well as proteins involved in SNARE and syntaxin-1 binding (e.g., Stxbp1, Syt1, Snap25, Snap91, Cplx1). These results further support the notion that genetic perturbations in PD converge on synaptic protein networks, marking early molecular signatures of dopamine dysfunction (Figure 5E).

### CRISPR screening identifies presynaptic modifiers of dopaminergic neuronal vulnerability

We hypothesized that alterations in the levels and/or distribution of Module 11 proteins could predispose dopaminergic neurons to disease-related stresses, including α-synuclein toxicity, aging, or environmental insults. To directly test our hypothesis, we conducted an in vivo cell type-specific CRISPR-based genetic screen to experimentally alter Module 11 proteins. Traditionally, cell survival CRISPR screens have been performed *in vitro* using cultured cell lines, which provide insights into basic cellular responses, but lack physiological context and cell-type specificity. To overcome these limitations, we leveraged a new CrAAVe-seq platform^33^, developed in the Kampmann lab (Figure 6A), to test the roles of these proteins in altering dopamine neuronal survival downstream of α-synuclein toxicity. This AAV-based pipeline enables selective recovery of sgRNA sequences from Cre-positive and Cre-negative cells via irreversible Cre-dependent recombination using distinct handle sequences (Figure 6B, Figure S6E).

We first optimized CrAAVe-seq delivery into dopamine neurons. Equal amounts of three AAVs were injected retro-orbitally, and we determined that 2.5 × 10^11^ genome copies (GC) achieved robust transduction of Cre-positive dopamine neurons while maintaining predominantly single-AAV infection, thereby minimizing confounding effects from multiple perturbations (Figure S6A–D). To estimate the effective library size under these conditions, we tested an oversized library containing 12,500 sgRNAs^33^ and found that ∼8,800 unique sgRNAs could be recovered from a single brain (Figure S6E–G). Based on these quantifications, we designed a pooled strategy to target 70 M11 genes, using 6 sgRNAs per gene. Five brains were pooled into each biological replicate, providing sufficient coverage to analyze ∼100 cells per sgRNA. In total, our library contained 467 sgRNAs (70 genes × 6 sgRNAs plus non-targeting controls, ∼10% of the total library). Guides were designed based on previously created mouse CRISPR knockout pooled library (Brie)^82^ and packaged into PHP.eB capsid AAV.

Pooled AAV-sgRNA libraries were delivered into Dat-Cre; LSL-Cas9 mice at 2 months of age, ensuring perturbations were restricted to dopamine neurons. After three weeks to allow for Cre-mediated recombination and target gene depletion, AAV encoding human α-synuclein harboring the pathogenic A53T mutation (A53T h-α-Syn) was stereotactically injected into the midbrain to induce partial dopamine neurodegeneration. Four weeks later, midbrain tissues were collected, and AAV episomes were isolated for quantification of sgRNA abundance via next-generation sequencing (NGS) (Figure 6C). By comparing gRNA representation between dopaminergic (Cre-positive) and control (Cre-negative) populations, we identified sgRNAs specifically depleted or enriched in dopaminergic neurons (Figure 6D).

To reliably identify genetic manipulations that modulate dopaminergic neuron vulnerability to α-synuclein-induced toxicity, we used MAGeCK^83^ to aggregate the effects of six sgRNAs per gene (Figure 6E). This approach ensured consistency across multiple sgRNAs targeting the same gene. This gene-level aggregation identified four presynaptic proteins, Stxbp1, Atp6v1b2, Atp6v1d, and Hspa8, whose depletion significantly exacerbated α-synuclein-induced neuronal vulnerability.

Among these hits, Stxbp1, involved in synaptic vesicle docking and neurotransmitter release, was the most significant when compared to control uninfected with A53T h-α-Syn. Additionally, Atp6v1b2 and Atp6v1d, integral components of the V-type ATPase complex, and Hspa8 were also identified as modulators whose depletion markedly exacerbated neuronal susceptibility. Remarkably, Hspa8 is a chaperone that in *Drosophila* is reported to suppress α-synuclein toxicity^84^, suggesting this function is conserved to mammals.

**Figure 6.**
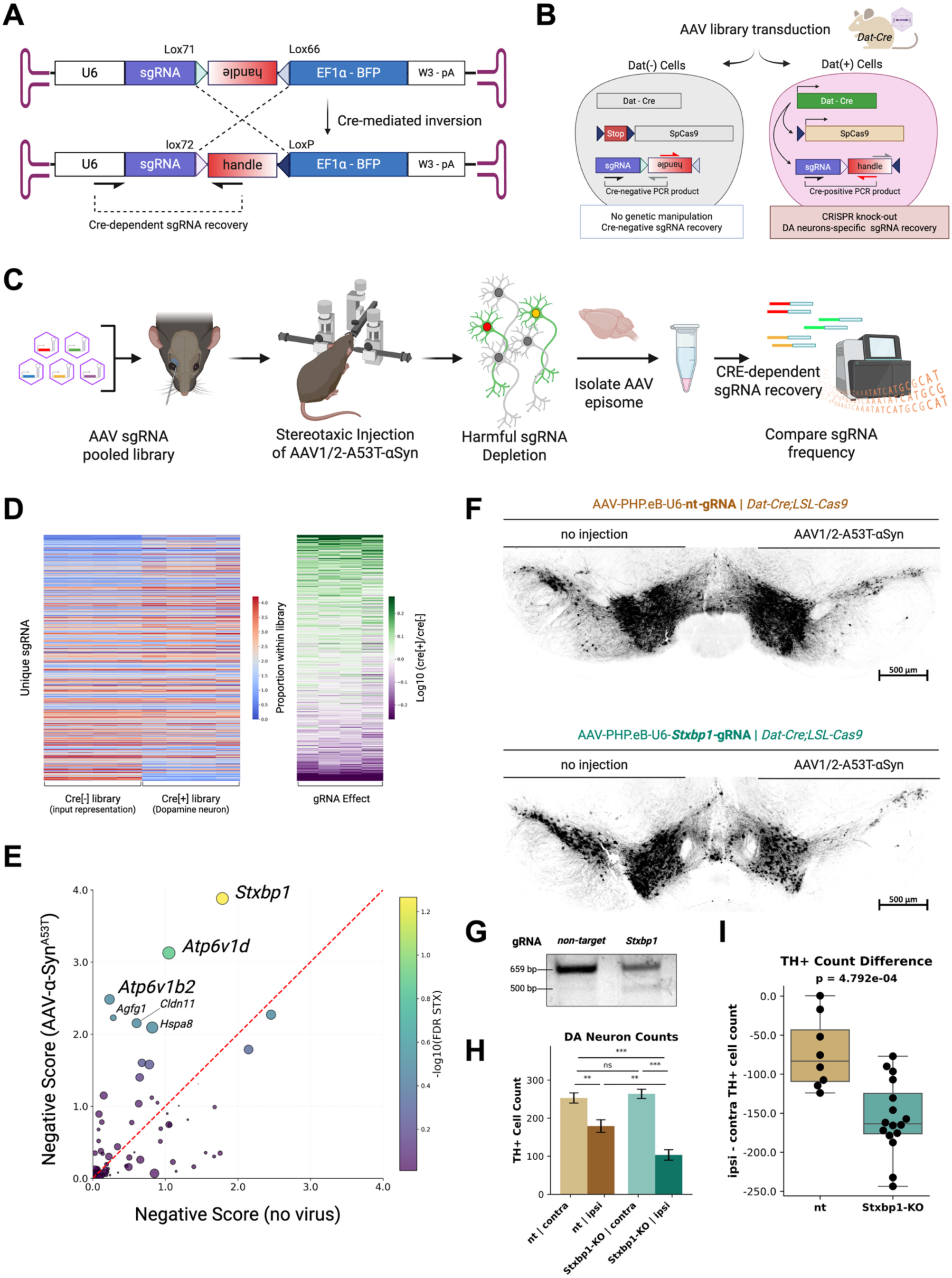
In Vivo CRISPR Survival Screening Identifies Stxbp1(Munc18-1) as a Protective Modifier Against α-synuclein Toxicity in Dopamine Neurons. (**A**) Schematic of the AAV construct enabling the Cre-dependent recovery of sgRNA sequences via irreversible handle recombination. (**B**) Strategy for simultaneous gene perturbation and recovery in dopaminergic neurons using Dat-Cre;LSL-Cas9 mice, enabling evaluation of gene function specifically within dopamine neurons. (**C**) Experimental timeline: pooled sgRNA library was delivered into 3–4-month-old Dat-Cre;LSL-Cas9 mice, followed by a four-week expression window before stereotactic injection of AAV encoding human A53T α-synuclein to induce toxic degeneration. Four weeks post-α-Syn injection, midbrain tissues were collected for episomal sgRNA recovery and next-generation sequencing (NGS). (**D**) Left panel: Quantification of sgRNA abundance by NGS showing relative representation of gRNAs in Cre-positive versus Cre-negative populations. Right panel: calculation of gRNA effect as the ratio of sgRNA representation between Cre-positive and Cre-negative neurons. (**E**) Scatter plot comparing MaGeCK-derived gene-level negative scores in the presence (vertical axis) and absence (horizontal axis) of α-synuclein overexpression. Genes in the upper left quadrant exhibit greater depletion under α-synuclein toxicity, suggesting a protective role. Marker size reflects the number of significant gRNAs per gene, and color indicates the FDR-adjusted p-value derived from MaGeCK analysis. (**F**) Representative coronal sections of the midbrain immunostained for TH, showing the contralateral and ipsilateral hemispheres in mice treated with either Stxbp1 or control gRNAs. (**G**) T7E1 assay in NIH3T3 cells expressing SpCas9 confirms the expected indel at the *Stxbp1* locus with the gRNA used for histological validation. (**H**) Bar plot showing the absolute TH-positive cell counts in the contralateral and ipsilateral hemispheres across both gRNA groups. Statistical comparison was performed using a linear mixed-effects model, with a significant interaction observed between gRNA and injection side (*β* = +86.6, *p* = 0.002), indicating that Stxbp1 loss exacerbated α-synuclein-induced dopaminergic neurodegeneration..Significance from paired t-test is indicated as follows:* p < 0.001, ** p < 0.01, * p < 0.05, ns p ≥ 0.05. (**I**) Box plot showing the difference in TH-positive cell counts between ipsilateral and contralateral hemispheres for each group (unpaired t-test, *p* = 4.792e-04; Control *n* = 8, Stxbp1 *n* = 16).

**Figure S6.**
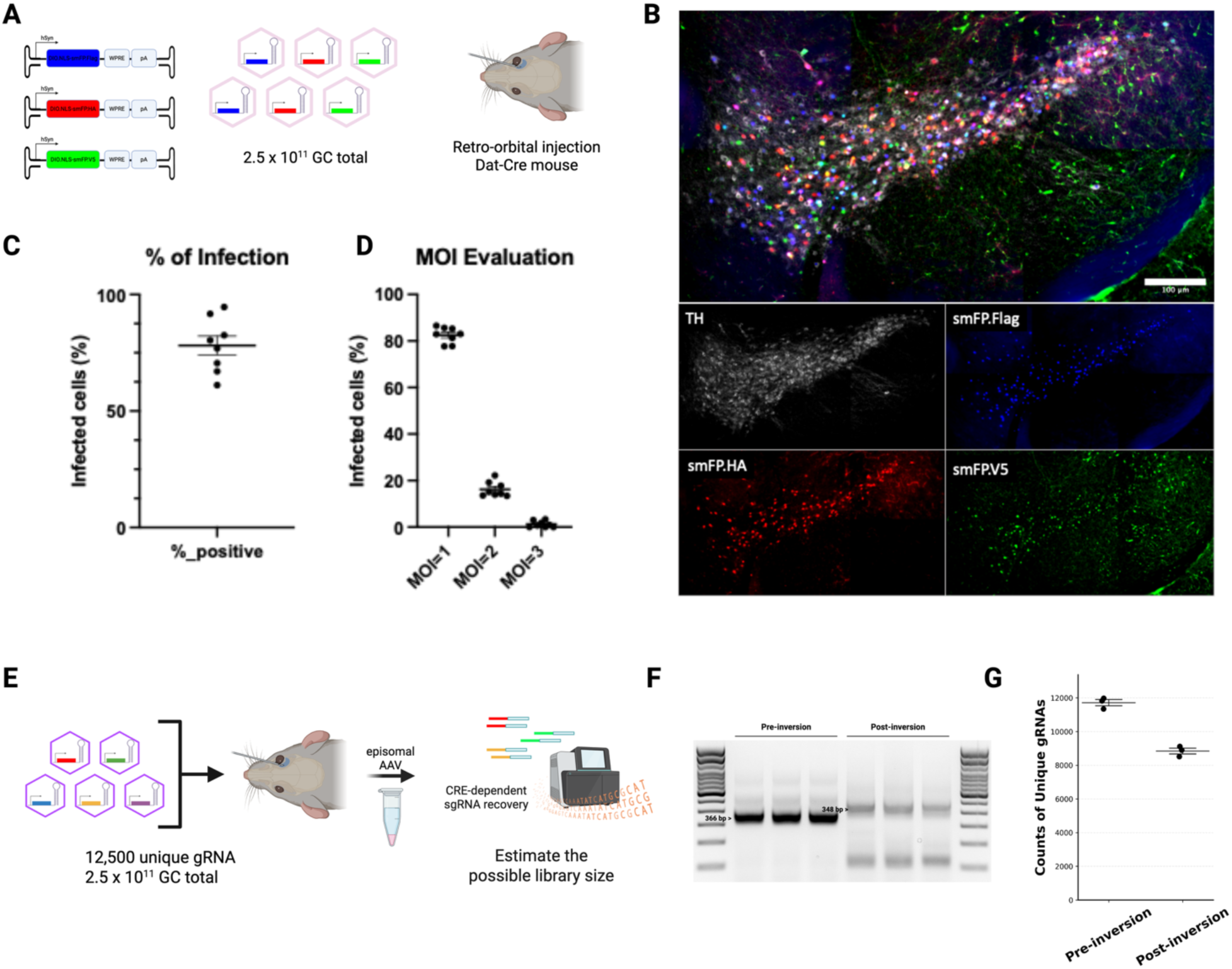
Optimization of CrAAVe-seq in dopamine neurons in vivo. (**A**) Estimation and optimization of viral titer for retro-orbital AAV transduction using equal proportions of epitope-tagged AAVs. (**B**) Representative coronal section of the midbrain showing dopaminergic neurons co-stained for TH, HA, V5, and Flag. (**C**) Quantification of transduction efficiency in dopamine neurons. (**D**) Evaluation of multiplicity of infection (MOI) based on co-staining patterns of epitope reporters. (**E**) Estimation of the achievable library size using an oversized test library of 12,500 unique gRNAs at a total dose of 2.5 × 10¹¹ GC. (**F**) Validation of Cre-dependent recombination enabling gRNA recovery from episomal AAV transgenes. (**G**) Quantification of unique gRNAs recovered pre- and post-inversion.

### Validation of Stxbp1/Munc18-1 as a Modifier of α-synuclein-Induced Neurodegeneration

To further validate the *in vivo* functional significance of the NGS gRNA candidate gene depletion, we performed targeted follow-up histological experiments focusing on *Stxbp1*, the most prominent hit from our screen. *Dat-Cre;LSL-Cas9* mice were retro-orbitally injected with AAVs delivering either Stxbp1-targeting or control gRNAs, followed by unilateral injection of AAV expressing human A53T α-Syn into the midbrain. TH-positive dopamine neurons were subsequently quantified in the midbrain region. In the non-target control gRNA, a significant reduction in TH-positive cells was observed in the ipsilateral hemisphere compared to the contralateral side, consistent with α-syn-induced neurotoxicity (paired t-test, t = –4.464, p = 0.0029, Cohen’s d = –1.58, n = 9 mice) (Figure 6F and G). Notably, mice receiving Stxbp1-targeting gRNAs exhibited even greater ipsilateral neuronal loss compared to controls, as reflected in the ipsi–contra difference (unpaired t-test, *t* = 13.225, *p* = 1.13 × 10⁻⁹, Cohen’s *d* = –3.31, n = 18 mice) (Figure 6H and I). To rigorously evaluate whether Stxbp1 depletion synergized with α-Syn toxicity, we employed a linear mixed-effects model incorporating both gRNA group and injection side as fixed effects while accounting for sample variability. This analysis revealed a significant interaction between *Stxbp1* deficiency and α-Syn injection (*β* = +86.6, *p* = 0.002), indicating that *Stxbp1* loss exacerbated α-Syn-mediated dopaminergic neurodegeneration beyond the effects of either manipulation alone (Figure 6H and I). Thus, the combined data indicate that *Stxbp1* is a critical presynaptic factor whose loss amplifies α-syn-induced vulnerability in dopaminergic neurons, reinforcing the concept that early synaptic dysfunction constitutes a key pathogenic mechanism in the prodromal period.

## Discussion

This study provides multi-dimensional evidence that disruptions in presynaptic vesicle trafficking constitute one of central and convergent pathogenic events at an early stage in mice downstream of mutations that in humans are associated with Parkinson’s disease (PD). By leveraging new tools, including in vivo proximity labeling (HiUGE-iBioID), dopamine neuron-specific subcellular proteomics (iBioCoFrac), and targeted functional validation through CRISPR screening, molecular alterations preceding overt dopaminergic neurodegeneration could be systematically delineated.

These methodologies synergistically reveal new insights into potential mechanisms that contribute to dopamine neuronal pathogenesis. It is essential, however, to acknowledge inherent limitations of the mouse models employed here with regard to human PD. While these genetic lines (*KI D620N Vps35*, *Tg-Th-hSnca^A30P/A53T^*, and *Tg-Pdgfb-hLRRK2^G2019S^*) model key genetic aspects of PD, they often present comparatively mild or late-onset neurodegeneration^64,73,76,85–89^. Nevertheless, their subtle phenotypes may effectively capture prodromal disease stages, offering potential insights into initial pathological processes that precede neuronal loss.

A central finding from our BioID analyses was the striking convergence of three pivotal PD-associated proteins: α-Syn, LRRK2, and VPS35, on presynaptic molecular machinery essential for synaptic vesicle docking, priming, fusion, and recycling. While previous studies have implicated broad vesicular and lysosomal dysfunction^90–93^ our data identified a presynaptic proteome of 74 proteins consistently within proximity to all three PD-related proteins. This set included proteins previously implicated in Parkinson’s pathobiology, such as Auxilin (Dnajc6) and Endophilin A1 (Sh3gl2). We observed significant enrichment for biological processes related to clathrin-mediated endocytosis, with strong associations in presynaptic compartments, exemplified by proteins such as Snap25, Snap91, Epn1, Epn15l1, and Clint1. Importantly, these findings represent the first unbiased in vivo mapping of a convergent molecular process shared by α-synuclein, VPS35, and LRRK2 under physiological expression conditions.

Mutant-specific BioID results further revealed distinctive remodeling of local proximity networks by individual PD mutations in vivo brain. The D620N mutation in VPS35 specifically enriched endocytic regulators such as Numb, Endophilin-B2, Smap1/2, and Fcho2, suggesting altered clathrin-coated pit regulation. The A30P mutation in α-synuclein increased associations with endosomal trafficking regulators including Rab11fip5, Acap2, Sgip1, and Annexin A6, alongside shifts in SNARE-related proteins such as Prrt2 and Complexin-1. In parallel, the G2019S mutation in LRRK2 showed neomorphic associations with Annexin A1, Epsin 2/3, and the Rab4/5 effectors Rabenosyn-5 and Rabaptin-5, as well as vesicle docking proteins Stxbp2 and Synaptotagmin-4. Together, these nuanced, genotype-specific alterations converge on the biological process of clathrin-mediated endocytosis and vesicle recycling, with a particular emphasis on the neuronal presynapse as the affected compartment.

Using dopamine neuron-specific iBioCoFrac, these proximity findings could be translated into cellular-level insights, uncovering consistent alterations in synaptic vesicle protein modules across three mouse models (LRRK2, SNCA, VPS35). Importantly, disruptions in synaptic vesicle regulation and clathrin-mediated endocytosis were observed in the prodromal period before reported dopamine pathology in each line, suggesting they may serve as early molecular signatures of dopamine neuronal defects. This spatial proteomics approach reinforced the presynapse as an early pathological epicenter, aligning molecular deficits closely with stages preceding dopamine neuronal loss.

Recent evidence highlights synaptic dysfunction (synaptopathy) as fundamental to PD pathogenesis, implicating presynaptic vesicle cycling, particularly endocytic retrieval and vesicle recycling as vulnerable processes^94^. PD-associated proteins profoundly disrupt these mechanisms. LRRK2 mutations, especially G2019S, impair synaptic vesicle endocytosis by phosphorylating key endocytic regulators such as endophilin-A and synaptojanin-1, slowing vesicle retrieval and recycling^91,95^ Pathologic α-synuclein aggregates inhibit SNARE complex formation necessary for exocytosis and directly interfere with clathrin-mediated endocytosis, precipitating severe disruptions in synaptic vesicle cycling^96–99^. Similarly, VPS35 mutations disrupt retromer-dependent endosomal recycling pathways critical for maintaining synaptic protein homeostasis, exacerbating α-synuclein accumulation and synaptic dysfunction^100–102^. Rab GTPases, integral to vesicular trafficking, also intersect significantly with these PD-related pathways, reinforcing a common theme of vesicle cycling impairment across both monogenic and sporadic PD cases^77,103,104^.

Critically, our CRISPR-based survival screening established direct causal links between presynaptic dysfunction and neuronal vulnerability. Among the presynaptic proteins identified, STXBP1 (Munc18-1), an essential SNARE-complex regulator crucial for vesicle docking and priming^105,106^, emerged as the most significant hit. The functional interaction between α-synuclein and the SNARE complex remains debated, with studies reporting both promotive and inhibitory effects on SNARE assembly^99,107–109^. One possible mechanism is that STXBP1/Munc18-1 may protect against pathological α-synuclein–induced excessive oligomerization of SNARE complexes^110^. While knockout of STXBP1/Munc18-1 leads to neuronal silencing^111^ previous work has shown that neuronal silencing by tetanus toxin does not cause dopaminergic neuron death^112^. However, it remains to be determined whether silencing increases neuronal vulnerability under pathological stress, such as excess α-synuclein. In addition, recent studies in cultured HEK293 cells and Drosophila suggest that STXBP1 may counteract α-synuclein aggregation through a non-canonical, chaperone-like function that prevents pathological deposition^113,114^. Consistent with this, decreased STXBP1 expression correlates with Lewy pathology in human PD brains^115,116^.

Additionally, our data implicate other presynaptic factors like V-ATPase subunits and HSPA8, crucial for vesicle re-acidification and clathrin uncoating, respectively. V-ATPase subunits are essential for maintaining proper vacuolar acidification, especially in synaptic vesicles and lysosomes, and disruptions in V-ATPase activity can impair synaptic vesicle recycling, lysosomal degradation, and proteostasis. Our iBioCoFrac analysis revealed altered abundance of specific V-ATPase subunits, supporting their early involvement in dopaminergic dysfunction. Recent studies have also highlighted TMEM175, a lysosomal proton channel, as a key regulator that functions closely with V-ATPase to maintain lysosomal pH. Dysfunction of TMEM175 exacerbates α-synuclein aggregation by disrupting lysosomal homeostasis, linking vesicle acidification failure to proteinopathy ^117^.

Similarly, HSPA8 (Hsc70), in conjunction with co-chaperones such as DNAJC6 (auxilin), mediates the critical step of clathrin coat removal from vesicles during endocytosis. In our BioID analysis, HSPA8 was detected in proximity to LRRK2, supporting its functional association with PD-linked trafficking networks; this is consistent with previous reports identifying HSPA8 and DNAJC6 in the endocytic axis affected by LRRK2 kinase activity^43,118^. Mutations in DNAJC6 are genetically linked to early-onset PD ^119^, supporting a role of HSPA8-mediated clathrin uncoating in maintaining synaptic vesicle recycling fidelity and neuronal survival. These proteins also significantly influenced neuronal susceptibility in our functional assays, reinforcing the critical importance of presynaptic trafficking integrity in PD. Future studies should further elucidate the molecular mechanisms and cooperative roles of these trafficking regulators within the broader presynaptic network to uncover novel intervention points for disease-modifying therapies.

In conclusion, the integrated, unbiased, and data-driven approach presented here reveals presynaptic vesicle cycling disruptions as one of central and causally significant events that precede dopamine neuronal loss downstream of mutations associated with human PD. The emergence of STXBP1 as a potential suppressor of dopamine vulnerability to α-synuclein, alongside broader synaptic trafficking proteins, provides an actionable framework for future research. Targeting presynaptic mechanisms to restore synaptic homeostasis may represent a highly promising strategy for halting or delaying PD-associated dopamine neuronal loss, particularly during its earliest, potentially reversible stages. Along these lines, genetic therapies, such as AAV-mediated gene delivery and antisense oligonucleotide-based modulation, are under active development for STXBP1-dependent neurodevelopmental disorders^120,121^. While more mechanistic studies are needed on the role of Stxbp1 in buffering a-syn toxicity, these approaches may offer readily adaptable future approaches.

## Acknowledgements

This research was funded in whole or in part by Aligning Science Across Parkinson’s (ASAP) P383000054 through the Michael J. Fox Foundation for Parkinson’s Research (MJFF). S.H.S was supported by NIH/NIMH MH126954. D.S. is supported by a PhRMA Foundation Predoctoral Fellowship in Drug Discovery. M.K. was supported by a Tau Consortium Investigator Grant from the Rainwater Charitable Foundation and by NIH/NIA grant R01 AG082141. Y.D was supported by NIH/NHGRI R35HG011328. We are grateful to Charles Gersbach, Ph.D. (Department of Biomedical Engineering, Duke University) for his consultation on the CRISPR screening analysis. We thank Nicole Calakos, M.D., Ph.D. and Elana Lockshin, Ph.D. for their kind instruction on the mouse surgical procedures. We also acknowledge Kathleen Miller, Omar Gammouh, and Matthew Trn for their general assistance with the research. We further acknowledge the Duke Proteomics and Metabolomics Shared Resource, the Duke Light Microscopy Core Facility, and the Duke Transgenic Core Facility for their technical support in completing this study.

## Author Contributions

S.S. conceptualized and supervised the study and contributed to manuscript writing. All authors reviewed and approved the final manuscript. D.S. conceived and designed the study, performed all experiments, data analyses, and figure preparation, and wrote the original manuscript. J.K., Y.G., and E.B. conducted mouse experiments and contributed to construct generation. A.O. optimized and performed the CRISPR screening methodology. Y.D. provided funding and partial methodological support for next-generation sequencing experiments. P.P. contributed to data analysis. E.S. performed and analyzed the proteomics experiments.

## Materials and Methods

### Animals

All animal procedures were performed under a protocol approved by the Duke University Institutional Animal Care and Use Committee in accordance with National Institutes of Health guidelines. Mice were group-housed in the Duke University Division of Laboratory Animal Resources facility under controlled conditions (ambient temperature of 72 ± 2°F, relative humidity of 30–70%, and a 12-hour light/dark cycle).

For the HiUGE iBioID experiment, three homozygous knock-in Parkinson’s disease (PD) mouse lines were employed. The LRRK2 G2019S knock-in line (strain: C57BL/6-Lrrk2tm4.1Arte, Taconic, Cat#13940), the Vps35 p.D620N knock-in line (JAX #023409), and an A30P SNCA line (generated at the Duke Transgenic Core) were each initially crossed with H11-Cas9 mice (JAX #28239) to generate double homozygous animals. These double homozygotes were subsequently mated with the respective homozygous PD lines so that all offspring were homozygous for the mutated PD protein and heterozygous for the Cas9 allele. H11-Cas9 mice, which are wild-type for the PD protein, served as controls.

For the iBioCOFrac experiment, PD model strains included Tg(SNCA A30PA53T)83Vle/J (JAX #008239), Tg(Lrrk2 G2019S)2Vlb/J (JAX #016575), and Vps35tm1Rck/J (JAX #023409), maintained as either hemizygous or heterozygous. These strains were crossed with B6.SJL-Slc6a3tm1.1(cre)Bkmn/J mice (JAX #006660) to generate double transgenic animals [Tg(SNCA A30PA53T); B6.SJL-Slc6a3tm1.1(cre)Bkmn/J, Tg(Lrrk2 G2019S); B6.SJL-Slc6a3tm1.1(cre)Bkmn/J, and Vps35tm1Rck/J; B6.SJL-Slc6a3tm1.1(cre)Bkmn/J]. The PD transgenic lines were then crossed with C57BL/6 wild-type females, yielding offspring with 50% PD transgenic; B6.SJL-Slc6a3tm1.1(cre)Bkmn/J (+/–) and 50% wild-type; B6.SJL-Slc6a3tm1.1(cre)Bkmn/J (+/–), with the non-transgenic progeny used as controls.

For the CRISPR screening experiment, LSL-Cas9 mice (JAX #024857) were crossed with B6.SJL-Slc6a3tm1.1(cre)Bkmn/J mice (JAX #006660) to generate double heterozygous animals for subsequent screening applications.

### HiUGE AAV plasmid preparation

Plasmid constructs for the HiUGE AAV system were generated following previously described methods^32,122^. In brief, each plasmid was engineered to include an HA-tagged TurboID coding sequence flanked by donor-specific guide RNA (DS-gRNA) recognition sites that are inert to the host genome.

For the Vps35 construct, the donor coding sequence was inserted at the extreme C-terminal end of the coding exon at the genomic Vps35 locus using a guide RNA with the sequence tctgaggggccaatctatga agg (GS-gRNA + PAM). In contrast, targeting the Snca locus posed challenges due to limited space at the C-terminal exon and the need to preserve its coding sequence. Therefore, an intronic targeting strategy was employed as previously described^32,123^. In this approach, the donor sequence encoding the HA-tagged TurboID was flanked by constitutive splicing acceptor and donor sequences and inserted into an intronic region using a guide RNA with the sequence aaggtggagtgttctaatgg ggg.

For the Lrrk2 construct, to avoid disrupting the C-terminal kinase domain, the HA-tagged TurboID was inserted into the N-terminal region. A custom donor construct was designed, comprising a self-cleaving T2A sequence, followed by a coding sequence from exon 1–2 of Lrrk2 fused in-frame to the HA-tagged TurboID. The T2A peptide mediates ribosomal skipping during translation, ensuring the production of N-terminal TurboID-tagged full length Lrrk2. This donor construct was also flanked by constitutive splicing acceptor and donor sequences to facilitate targeted insertion into intron 2 of the Lrrk2 locus. The guide RNA used for Lrrk2 targeting was ctgctaaattatcaaaccag agg.

To enable endogenous expression of soluble TurboID for background detection, an additional donor construct was designed. In this construct, sequences at the C-terminal region or 3′-UTR were targeted to introduce a stop codon followed by an internal ribosome entry site (IRES) and the TurboID-HA coding sequence.

### AAV preparation

AAV were prepared following previously described methods^32,34^. For large-scale production intended for in vivo applications, HEK293T cells were cultured in six 15-cm dishes. Cells were transfected via a triple-transfection protocol using 15 μg of the HiUGE vector, 30 μg of pADdeltaF6, and 15 μg of the serotype plasmid (pUCmini-iCAP-PHP.eB, Addgene plasmid #103005). Three days post-transfection, cells were lysed and the virus was purified and concentrated using an Optiprep density gradient (Sigma #D1556). The concentrated virus was aliquoted and stored at – 80°C until further use.

For small-scale production intended for in vitro cell culture, HEK293T cells were grown in 12-well plates and transfected using the same triple-transfection strategy with 0.4 μg of the HiUGE vector, 0.8 μg of pADdeltaF6, and 0.4 μg of the serotype plasmid (pUCmini-iCAP-PHP.eB, Addgene plasmid #103005). Three days post-transfection, the virus-containing medium was collected and filtered through Costar Spin-X columns (Sigma #8162). For each preparation, 0.5 mL of AAV-containing supernatant was filtered and stored at 4°C until use.

### AAV-mediated HiUGE-iBioID sample preparation

The HiUGE-iBioID sample preparation was performed based on previously described methods^32^ with minor modifications tailored to each target protein. For Snca and Vps35 tagging, neonatal (P0–2) H11-Cas9 mice were anesthetized by hypothermia and injected intracranially with purified AAV (PHP.eB serotype, titer >10^10 GC/μL) at a volume of 2 μL per hemisphere. To achieve optimal AAV transduction and HiUGE integration efficiency for Lrrk2 tagging, E15.5 embryonic fetuses were injected in utero with 1 μL per hemisphere of the same AAV preparation. IRES-TurboID constructs were used as negative controls.

At three weeks of age, all pups received daily intraperitoneal injections of biotin (50 mg/kg) for five consecutive days. Mice were then deeply anesthetized with isoflurane and euthanized by decapitation. Brain tissue spanning from the forebrain to the midbrain was collected one day after the final biotin injection, following removal of the cerebellum and pons, and immediately snap-frozen at –80 °C until further processing.

For protein purification, brain tissues from three mice (for Vps35 and Snca samples) or five mice (for Lrrk2 samples) were pooled. The combined tissues were homogenized and sonicated in RIPA lysis buffer supplemented with cOmplete protease inhibitor cocktail (Sigma, Cat#11873580001), and then centrifuged at 16,000 × g for 30 minutes at 4 °C. The supernatant lysates were normalized based on protein concentration across samples and desalted using Zebra columns with a 7 K MWCO (ThermoFisher, Cat#89894 or Cat#89892). The flow-through was incubated with 50 μL magnetic Strepavidin beads (Pierce, Cat#88816) at 4 °C overnight. The beads were subsequently washed sequentially as follows: twice with RIPA buffer, once with 1 M KCl, once with 0.1 M Na₂CO₃, once with 2 M urea in 10 mM Tris-HCl, and twice with RIPA buffer. Finally, biotinylated proteins were eluted by boiling the beads in 90 μL elution buffer (2% SDS, 25 mM Tris, 10 mM DTT, and 2.5 mM biotin) and used for downstream LC-MS/MS and Western blot analyses.

### HiUGE-iBIoID LC-MS/MS data acquisition and analysis

Sample Preparation and LC-MS/MS Analysis; BioID samples (three bait–TurboID fusion and three soluble–TurboID control) were processed by the Duke Proteomics and Metabolomics Core Facility (DPMCF). Samples, stored at −80 °C until processing, were spiked with 1–2 pmol undigested bovine casein as an internal quality control standard, reduced for 15 min at 80 °C, and alkylated with 20 mM iodoacetamide for 30 min at room temperature. After addition of 1.2% phosphoric acid and 733 µL of S-Trap binding buffer (90% MeOH/100 mM TEAB), proteins were trapped on S-Trap cartridges, digested with 20 ng/µL trypsin (Promega) for 1 h at 47 °C, and sequentially eluted with 50 mM TEAB, 0.2% FA, and 50% acetonitrile/0.2% FA. Samples were lyophilized, resuspended in 48 µL of 1% TFA/2% acetonitrile containing 12.5 fmol/µL yeast ADH, and pooled to generate a study pool QC (SPQC). For LC-MS/MS, 2 µL of each sample was analyzed on an MClass UPLC system (Waters Corp.) coupled to a Thermo Orbitrap Fusion Lumos mass spectrometer with a FAIMSPro device and nanoelectrospray ionization. Peptides were trapped on a Symmetry C18 column (20 mm × 180 µm, 5 µL/min, 99.9/0.1 water/acetonitrile) and separated on a 1.8 µm Acquity HSS T3 C18 analytical column (75 µm × 250 mm) with a 90-min gradient from 5% to 30% acetonitrile/0.1% formic acid at 400 nL/min and 55 °C. Data collection was performed across compensation voltages of −40 V, −60 V, and −80 V in data-dependent acquisition (DDA) mode, with full MS scans (m/z 375–1500, r = 120,000 at m/z 200, AGC target 4e5) followed by MS/MS scans acquired in the linear ion trap under HCD (30%) in rapid mode (AGC target 1e4, maximum fill time 35 ms). Each CV cycle was 0.66 s, with 2 s between full MS scans, and a 20 s dynamic exclusion was applied. The total run time per injection was approximately 2 h.

Quantitative Data Analysis; A total of 12 UPLC-MS/MS analyses were performed (excluding conditioning runs but including three replicate SPQC samples). Data were imported into Proteome Discoverer 3.0 (Thermo Scientific Inc.) and aligned using the Minora Feature Detector algorithm based on accurate mass and retention time of precursor ions. Relative peptide abundance was determined from ion chromatogram peak intensities across aligned features. MS/MS data were searched against the SwissProt Mus musculus database (downloaded August 2022), a contaminant/spike-in database (bovine albumin, bovine casein, yeast ADH), and reversed-sequence decoys for false discovery rate estimation. Sequest with INFERYS was used for fragment ion prediction and database searching. Parameters included fixed carbamidomethylation on Cys, variable oxidation on Met, precursor mass tolerance of 2 ppm, product ion tolerance of 0.8 Da, and full trypsin specificity. Peptide and Protein FDR Validator nodes applied a maximum 1% protein-level false discovery rate. Peptide homology was resolved using razor rules, assigning shared peptides to the protein with the most identified peptides. Protein groups with identical peptide sets were consolidated, and a master protein was assigned based on sequence coverage.

### Statistical Analysis of Proximity Proteome Overlap

To evaluate the significance of the overlap among the two or three proximity proteomes, we applied the hypergeometric test, which determines whether the observed intersection exceeds random expectation based on the total detected proteins. For each pairwise comparison and the three-way overlap, we computed the probability of observing at least the given number of shared proteins using a hypergeometric distribution, implemented through the survival function (scipy.stats.hypergeom.sf). This analysis was conducted independently for each dataset combination, ensuring that statistical significance was not affected by the order of set selection.

### iBioCoFrac Proteomics Sample Preparation

The Cre-dependent TurboID construct was generated by cloning the TurboID coding sequence from V5-TurboID-NES_pCDNA3 (Addgene #107169) into a pAAV-hSyn-DIO vector, where “DIO” denotes a double-floxed inverted open reading frame that permits Cre-mediated recombination. AAV-PHP.eB capsids packaging the hSyn-DIO-TurboID construct were injected via E15.5 in utero fetal brain injection into all pups within both uterine horns. Genotyping was performed postnatally to distinguish PD transgenic from wild-type offspring.

At 6–7 weeks of age, both PD transgenic and littermate wild-type animals, all carrying Dat-Cre heterozygously, received daily intraperitoneal injections of biotin (50 mg/kg) for five consecutive days. Mice were then deeply anesthetized with isoflurane and euthanized by decapitation. Subcortical and midbrain regions were dissected by removing the cortex, olfactory bulb, cerebellum, and pons, thereby enriching for dopamine neuron populations spanning the basal ganglia (including the striatum) and the dopaminergic midbrain nuclei (SN/VTA). Freshly dissected brain samples from five brains of the same genotype (sex-mixed) were pooled to constitute one replicate. In each experiment, one replicate was prepared for PD transgenic animals (n = 5) and one replicate for wild-type animals (n = 5). Three replicates per strain were performed. The pooled samples were subsequently processed using LOPIT-DC subcellular fractionation for downstream proteomic preparation.

### iBioCoFrac Dopamine Neuron Enriched Subcellular Fractionation

Pooled brain samples were processed using LOPIT-DC subcellular fractionation as described previously^68^. Three independent fractionation experiments were performed per strain. Freshly dissected brains were immediately placed in a 2 mL Dounce homogenizer on ice containing 1 mL of isotonic TEVP homogenization buffer (320 mM sucrose, 10 mM Tris base, 1 mM EDTA, 1 mM EGTA, 5 mM NaF, pH 7.4), supplemented with cOmplete protease inhibitor cocktail (Sigma, Cat#11873580001). Tissues were homogenized with 15 passes using the Dounce homogenizer, and the lysate was subsequently brought to a final volume of 5 mL with additional TEVP buffer. To further release organelles, the lysate was passed through a 0.5 mL ball-bearing homogenizer (14 mm ball, Isobiotec) for two passes, yielding a final lysate volume of approximately 7.5 mL per sample.

The lysates were aliquoted into replicate microfuge tubes (Beckman Coulter, Cat#357448) and subjected to differential centrifugation following Geladaki et al.’s LOPIT-DC protocol^67^. Centrifugation was performed at 4 °C in a tabletop Eppendorf 5424 centrifuge at sequential speeds of 200 g, 1000 g, 3000 g, 5000 g, 9000 g, 12,000 g, and 15,000 g. To isolate the final three fractions, an ultracentrifugation step was conducted at 4 °C using a Beckman TLA-100 ultracentrifuge with a TLA-55 rotor at speeds of 30,000 g, 79,000 g, and 120,000 g, respectively. Throughout the process, samples were kept on ice and pellets were stored at –80 °C. Pellets from seven fractions (those corresponding to spins from 5000 g to 120,000 g) were carried forward for further processing.

Each pellet was resuspended in 500 µL of RIPA buffer supplemented with cOmplete protease inhibitor, then sonicated. The samples were subsequently centrifuged at 15,000 rpm for 30 minutes at 4 °C. Following normalization of protein concentrations across the seven fractions for each genotype, 20 µL of streptavidin magnetic beads were added per sample. Samples were rotated at 4 °C overnight to capture biotinylated proteins. The beads were then washed following the protocol established for the HiUGE-iBioID sample preparation: twice with RIPA buffer, once with 1 M KCl, once with 0.1 M Na₂CO₃, once with 2 M urea in 10 mM Tris-HCl, and finally twice with RIPA buffer. Finally, biotinylated proteins were eluted by boiling the beads in 50 µL of elution buffer (2% SDS, 25 mM Tris, 10 mM DTT, and 2.5 mM biotin) and used for downstream TMT multiplexing and Western blot analysis.

### iBioCoFrac TMT-multiplexed quantitative LC-MS/MS analysis

Sample Preparation; Protein compositions were examined from centrifuged fractions (F4–F10) of three independent study groups, each including wild-type (WT) and transgenic (Tg) samples enriched with biotin. In total, 42 samples (3 replicates each of WT_F4–WT_F10 and Tg_F4–Tg_F10) were processed at the Duke Proteomics and Metabolomics Core Facility (DPMCF). Samples were stored at −80 °C until processing, supplemented with 3% SDS, and spiked with 1–2 pmol casein as an internal quality control standard. Reduction was performed with 10 mM dithiothreitol for 30 min at 32 °C, followed by alkylation with 20 mM iodoacetamide for 45 min at room temperature. Proteins were acidified with 1.2% phosphoric acid and combined with 437 µL of S-Trap binding buffer (90% methanol/100 mM TEAB), trapped on S-Trap microcartridges, digested using 20 ng/µL sequencing grade trypsin (Promega) for 1 h at 47 °C, and sequentially eluted with 50 mM TEAB, 0.2% formic acid, and 50% acetonitrile/0.2% formic acid. All eluates were lyophilized to dryness.

Offline Fractionation and LC-MS/MS Analysis; Dried peptides were fractionated using the Pierce™ High pH Reversed-Phase Peptide Fractionation Kit (ThermoFisher Scientific), with fractions 1 and 7 as well as fractions 2 and 8 combined, yielding six fractions per sample set. After lyophilization, 3 µL (25%) of each fraction was analyzed using an MClass UPLC system (Waters Corp.) coupled to a Thermo Fusion Lumos mass spectrometer equipped with a FAIMS ion-mobility device and nanoelectrospray ionization source. Peptides were first trapped on a Symmetry C18 column (20 mm × 180 µm, 5 µL/min, 99.9/0.1 v/v water/acetonitrile) and then separated on a 1.8 µm Acquity HSS T3 C18 analytical column (75 µm × 250 mm) with a 90-min linear gradient of 5–30% acetonitrile/0.1% formic acid at 400 nL/min and 55 °C. Data collection was performed in data-dependent acquisition (DDA) mode with full MS scans (m/z 375–1575, r = 120,000 at m/z 200) across compensation voltages of −40 V, −60 V, and −80 V. MS/MS scans were acquired at r = 50,000 with an AGC target of 200% and a 120 ms maximum injection time. Dynamic exclusion was set at 45 s, and total run time per injection was approximately 2 h.

Data Processing and Quantitative Analysis; MS data were processed in Proteome Discoverer 3.0 (Thermo Scientific Inc.) for extraction of TMT16 reporter ion signals (m/z 126, 127N, 127C, etc.). Database searches were performed using the Sequest algorithm against the SwissProt Mus musculus database (downloaded August 2022), a common contaminant/spike-in database (e.g., bovine albumin, bovine casein, yeast ADH), and reversed-sequence decoys for false discovery rate (FDR) estimation. Search parameters included fixed modifications on Cys (carbamidomethylation), Lys (TMT), and peptide N-termini (TMT), with variable modification on Met (oxidation). Mass tolerances were 2.0 ppm for precursors and 0.02 Da for product ions, with full trypsin specificity required. A 1% protein-level FDR cutoff was applied using Peptide Validator and Protein FDR Validator nodes. Peptide homology was resolved using razor rules, assigning shared peptides to the protein with the greatest peptide evidence, and protein groups were defined by shared peptide sets, with master proteins designated based on percent sequence coverage.

### Principal Component Analysis

To assess whether the proteomic profiles across the fractionated samples (fractions 4–10) exhibited distinct segregation, Principal Component Analysis (PCA) was conducted. The dataset included seven fractions, each with three independent biological replicates, as well as six standardized pooled quality control (SPQC) samples, totaling 27 data points.

PCA was performed using the scikit-learn Python package, applied to z-score normalized spectral count data. The data was standardized by mean-centering and scaling to unit variance, and the principal components were computed using Singular Value Decomposition (SVD) from the covariance matrix. The first four principal components (PCs) were retained, and the variance explained by each PC was calculated to evaluate their relative contributions.

The PCA results were visualized in a 2D scatter plot based on PC1 and PC2. To highlight distribution trends, confidence ellipses were constructed using the covariance matrix of PC1 and PC2, with their major and minor axes scaled to a 95% confidence interval (*χ* 2 value = 5.991 for 2 degrees of freedom).

### Protein-level Comparison of the iBioCoFrac data

For protein-level analysis of the iBioCoFrac data, we performed a two-way repeated measures ANOVA to examine the effects of genotype (wild-type vs. transgenic) and biofraction (F4-F10) on relative protein intensities across seven biofractions. The analysis was conducted using the statsmodels AnovaRM function in Python, with mixture as the subject variable and genotype and biofraction as within-subject factors. Only genes meeting specific criteria were included in the analysis: exactly 2 genotypes, 3 biological replicates, and 7 biofractions per gene. For each qualifying gene, we calculated F-values, degrees of freedom, and p-values for main effects (genotype, biofraction) and their interaction (genotype × biofraction). To assess the biological significance of observed differences, we computed Cohen’s d effect sizes for each biofraction by comparing mean relative intensities between genotypes, using the pooled standard deviation as the denominator. We then calculated composite scores by multiplying the negative log10-transformed p-values by the maximum effect size across biofractions, enabling ranking of genes by both statistical significance and biological effect magnitude.

### Module detection

Co-fractionated proteins were clustered following the method described previously^68^. Briefly, an enhanced adjacency matrix derived from the protein covariation data was constructed and then clustered in Python using the Leiden algorithm^78^, an improvement upon the Louvain algorithm. The Leiden algorithm optimizes the partitioning of a graph into modules by maximizing a quality statistic. In this study, we employed the ‘Surprise’ quality statistic^78^ to determine the optimal modular partitions of the protein covariation graph.

### Module-level Significance Evaluation

To evaluate clusters that had a significant change in protein distributions between WildType and Transgenic genotypes, we extended MSstatsTMT Linear Mixed-Effects Modeling framework. We aggregated proteins across mixtures by sum-normalizing the protein intensities within each protein (relative intensity)-- the variance between mixtures is negligible after normalization. We defined the response variable as the log2-transformed relative protein intensity. For each cluster, the subset of proteins assigned to that partition was fit with a mixed-effects model treating Protein as a random effect, to account for baseline variation among proteins, and Condition as a fixed effect:

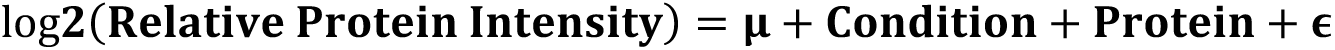

The response variable is defined as the log₂-transformed, sum-normalized protein intensity for all proteins within a cluster. To evaluate genotype effects at the module level, we constructed a contrast vector specifying the difference between the Transgenic and Wild-Type fixed effects (TG–WT). This contrast was assessed using the lmerTest package in R, yielding log₂ fold-change estimates, standard errors, test statistics, and raw p-values per cluster. We control for multiple testing across modules with Bonferroni correction; modules with adjusted p-values < 0.05 were considered significant (Vps35: n = 17, Lrrk2: n = 42, Snca: n = 17).

### Comparative analysis of HiUGE-iBioID and iBioCoFrac

To contextualize our HiUGE-BioID-based discovery of differential protein-protein interactions associated with Parkinson’s disease (PD) mutants, we performed a comparative analysis with subcellular co-fractionation proteomics data obtained from three PD models: transgenic α-SynA30P*A53T, LRRK2G2019S, and KI-D620N-Vps35. Our iBioCoFrac experiments and subsequent bioinformatic analyses identified co-varying protein modules in dopamine neurons, revealing 12 modules for VPS35, 15 modules for LRRK2, and 10 modules for α-synuclein.

To determine whether the observed altered spatial distribution of proteomes reflects impaired protein-protein interactions due to these PD-associated missense mutations, we applied a statistical hypergeometric test. This analysis evaluated the significance of the overlap between the spatial proteomics datasets and the protein-protein interaction (PPI) networks curated for α-synuclein, LRRK2, and VPS35. The goal was to elucidate the molecular underpinnings of dysregulated spatial modules within the PD protein interactome and to pinpoint key proteins directly linked to PD-related abnormalities arising from point mutations.

### iBioCoFrac Modulized Protein network visualization

Protein networks were visualized in Cytoscape (version 3.10.3) using data derived from a network-enhanced (NE)^124^ covariance-clustering matrix generated in the iBioCoFrac experiment, retaining only high-confidence covariation (NE score > 1.00). Filtered covariation data were imported into Cytoscape, where the Prefuse Force Directed Layout algorithm was applied with NE scores as heuristic edge weights, a default spring coefficient and length, and 100 correlation-based connections. Weak edges below the NE threshold were excluded to highlight significant covariation patterns and improve network clarity. Nodes were color-coded discretely according to their assigned module, while Log2FC and Log10_Padjust values were mapped to continuous color gradients for comparative analysis.

### Gene set enrichment analysis

Gene ontology (GO) enrichment analysis was performed on the entire proteomics dataset against a custom statistical domain comprising all identified brain proteins (10,112 unique proteins; Supplementary Data) from cumulative mouse brain proteomic studies in our lab (n = NNN) using ShinyGO version 0.80. All three GO categories—Biological Process, Molecular Function, and Cellular Component—were analyzed. For the Molecular Function category, a pathway size boundary of 10 to 500 was applied to exclude ambiguous terms, while default pathway size boundaries were used for the other queries. An FDR cutoff of 0.05 was used to determine significance.

### sgRNA library cloning for AAV screening

Seventy genes within module 11 were selected, and for each gene, six unique guide RNAs (gRNAs) were retrieved from the publicly available Mouse CRISPR Knockout Pooled Library (Brie; Pooled Libraries #73632, #73633, #73633-LV). The pooled oligonucleotides were synthesized by Twist Bioscience and subsequently cloned into the AAV vector pAP215 (Kampman lab) using inFusion cloning with BstXI and BlpI.

The ligated plasmid library was transformed into Endura ElectroCompetent Cells (≥1 × 10^10 cfu/µg DNA; VWR, Cat#71003-038 [Supplier No.60242-2]) by electroporation using a 0.2 cm cuvette with settings of 25 µF, 200 Ω, and 3000 V. Following electroporation, cells were recovered at 37°C for 1 hour in recovery medium, then cultured in 500 mL LB supplemented with ampicillin for 12–14 hours under shaking conditions until an OD of 0.07 was reached. Ten colonies were plated on LB-ampicillin plates and screened by colony PCR to verify unique insertion of the gRNA in the desired region of the plasmid vector.

Maxiprep purification of the library yielded 15 µg of plasmid DNA, which was used for AAV production. For AAV preparation, the library plasmid was co-transfected into HEK293T cells along with 30 µg of pADdeltaF6 and 15 µg of the serotype plasmid pUCmini-iCAP-PHP.eB (Addgene #103005), following the protocol described in the AAV preparation section. Three days post-transfection, cells were lysed, and the virus was purified and concentrated using an Optiprep density gradient (Sigma, Cat#D1556). The concentrated virus was aliquoted, normalized to 1.0 × 10^10 GC/μL, and stored at –80°C until further use.

### Library AAV injection

Two-month-old LSL-Cas9;Dat-Cre mice were used for library AAV injections. Mice were placed in an induction chamber and anesthetized with 3–4% isoflurane delivered at 0.8 LPM via a calibrated isoflurane vaporizer. Anesthesia was maintained until mice showed no response to foot pressure. Using an insulin syringe fitted with a 28-gauge needle, 100 µL (0.1 mL) of non-replicating adeno-associated virus (AAV) library—normalized to a concentration of 1.0 × 10^10 GC/μL—was administered.

### Stereotaxic Injection of AAV-Encoding Human A53T α-synuclein

Adult mice were initially anesthetized in an induction chamber using 3–4% isoflurane delivered at 0.8 LPM. Anesthesia was maintained at 1.5–2% during surgery, with the animal’s responsiveness (assessed via foot pressure every 10 minutes) ensuring adequate depth. Following induction, mice were secured in a stereotaxic frame after scalp shaving, ophthalmic ointment application, and disinfection of the surgical area with sterile alcohol and betadine. Supplemental heat was provided throughout the procedure via a heat pad covered in sterile drapes.

Using a Nanoject Nano-injector, 3.0 × 10^9^ genome copies (GC) in 600 nL of concentrated AAV1/2-CMV/CBA-human-A53T α-synuclein (Charles River Laboratories, GD1001-RV) was administered into the substantia nigra pars compacta (SNpc) at coordinates of AP –3.0 mm, ML +1.2 mm, and DV +4.3 mm from bregma. Injections were performed either unilaterally or bilaterally, depending on the experimental design.

### sgRNA recovery and sequencing for CrAAVe-seq

CrAAVe-seq sample preparation was performed following the protocol described in Ramani et al., 2025 with modifications^33^. Briefly, each brain was placed in a Dounce homogenizer containing TRIzol reagent (Thermo Fisher Scientific, 15596026) and thoroughly homogenized. Chloroform was added, and the mixture was vigorously shaken and centrifuged at 12,000 × g for 15 min at 4°C to separate the aqueous and organic phases. The aqueous phase (top layer) was collected, and nucleic acids were precipitated by adding isopropanol, and centrifuging at 12,000 × g for 10 min at 4°C. The pellet was washed with 75% ethanol in DNase/RNase-free water, centrifuged again, air-dried, and resuspended in DNase/RNase-free water (Thermo Fisher Scientific, AM9937). To remove residual RNA, the sample was treated with RNase A (Thermo Fisher Scientific, EN0531) at 37°C for overnight with gentle agitation. The resulting episomal DNA was either purified using a Zymo DNA Clean & Concentrator Kit (Zymo Research, D4013) and eluted in DNase/RNase-free water or stored at −20°C without column purification.

Episomal DNA was subjected to two distinct PCR amplification strategies to distinguish sgRNAs expressed in Cre-positive versus Cre-negative cells. The CrAAVe-seq plasmid pAP215 was designed with a Lox71/Lox66-flanked "handle" sequence, which undergoes unidirectional inversion in cells expressing Cre recombinase, thereby enabling cell type-specific sgRNA enrichment.

For Cre-inverted sgRNA amplification, we used a common forward primer (5’-GGCTTAATGTGCGATAAAAGACAG-3’) and a reverse primer specific to the Cre-inverted handle (5’-TGACTGGTACTGACACGTCG-3’). PCR was performed using Phusion High-Fidelity DNA Polymerase (Thermo Fisher Scientific, F530S) with 31 cycles, an annealing temperature of 55°C, and an extension step at 72°C for 15 sec. To amplify all sgRNAs present in the recovered episomes—regardless of Cre-dependent inversion—a common forward primer (5’-GGCTTAATGTGCGATAAAAGACAG-3’) was paired with a reverse primer independent of the Cre handle (5’-CGACTCGGTGCCACTTTTTCAAG-3’). PCR was performed with 20 cycles, an annealing temperature of 55°C, and an extension step at 72°C for 15 sec. Both amplified products were purified using SPRISelect beads (Beckman Coulter, B23317) following the manufacturer’s protocol and eluted in 100 µL DNase/RNase-free water.

### Next-Generation Sequencing Sequencing

Following DNA isolation from pre and post samples as described above, we prepared the samples for NGS sequencing and quantification as described below. Firstly, a primer extension step was done using the following primers for pre (ACACTCTTTCCCTACACGACGCTCTTCCGATCTNNNNNNNNNNCGACTCGGTGCCACTTTTTC AAG) and post (ACACTCTTTCCCTACACGACGCTCTTCCGATCTNNNNNNNNNNTGACTGGTACTGACACGTCG) samples. The ‘N’s in the primer sequence represented random unique molecule identifier (UMI) sequence. The primer extension reaction was prepared by mixing: 13ul of DNA sample, 10µl Q5 reaction buffer, 1µl of 10mM dNTP mixture, 1µl of 10µM UMI-containing primers, 1µl of Q5 polymerase, and 23µl of ultra-pure molecular biology water (Genesee Scientific, 18-193). The following thermocycler settings were used: an initial denaturation at 98°C for 1 minute, annealing at 55°C for 30 seconds, elongation at 72°C for 20 seconds, and a final 4°C hold. Following primer extension, the DNA was cleaned using SPRIselect beads (Beckman Coulter, B23319) following manufacturer’s protocol and using a 0.95:1 beads-to-sample volume ratio. After this, a pre-sequencing PCR was carried out as outlined below. Firstly, a PCR reaction was prepared by mixing the following: The primer extension reaction was prepared by mixing: 18ul of DNA sample (from the primer extension reaction), 5µl Q5 reaction buffer, 0.5µl of 10mM dNTP mixture, 0.5µl of 10µM common i7 primer (that is, common to both pre and post samples) (primer sequence: CAAGCAGAAGACGGCATACGAGATGTCTCGTGGGCTCGGAGATGTGTATAAGAGACA GTGAGACTATAAATATCCCTTGG), 0.4µl of 10µM indexed Truseq i5 primer (AATGATACGGCGACCACCGAGATCTACAC[10bp_index]ACACTCTTTCCCTACACGAC), and 0.8µl of Q5 polymerase. The following thermocycler setting was used for the PCR reaction: an initial denaturation for 30 seconds at 98°C; 14 cycles of denaturation (98°C, 10 seconds), primer annealing (48°C, 30 seconds), and elongation (72°C, 20 seconds); followed by a one-time final elongation at 72°C for 5 minutes and a 4°C hold. Following this PCR reaction, DNA were cleaned using SPRIselect beads as previously described. An extra PCR was done to add an i7 sample index for the purpose of nucleotide balancing during sequencing. This final PCR reaction was similar to the previous PCR reaction, only with the exception of using the indexed Nextera i7 primer as the forward primer (CAAGCAGAAGACGGCATACGAGAT[10bp_index]CTCGTGGGCTCGGAGATGTGTATAAGAGACAG), and the P5 sequence (AATGATACGGCGACCACCGAGATCTACAC) as the reverse primer—for both, 0.5µl of a 10µM primer solution was used. For this final PCR, the following thermocycler setting was used: an initial denaturation for 30 seconds at 98°C; 13 cycles of denaturation (98°C, 10 seconds), primer annealing (58°C, 30 seconds), and elongation (72°C, 20 seconds); followed by a one-time final elongation at 72°C for 5 minutes and a 4°C hold. Following the final PCR, DNA amplicons were purified using agarose gel electrophoresis to remove primer dimers and non-specific fragments before sequencing. All primers were purified using denaturing polyacrylamide gel electrophoresis.

### CRISPR Screen Analysis

Fastq files were demultiplexed using cutadapt (version 4.9), with selection for reads that contain the correct plasmid sequence, for pre and post samples. Using umitools (version 1.1.5), we named each read using the unique UMI sequence and discarded plasmid sequences. Using the index function in bwa (version 0.7.18), the guide RNA sequences were used to make a custom reference file with which the plasmid-derived reads were aligned. Next, taking only primary alignments, the aligned reads of all detected unique guide RNAs were counted for both pre and post reads. The code used for this analysis can be found here: https://github.com/Arinze-BioX/daichi_etal_parkinsons.

To analyze sgRNA representation, total count normalization was performed by dividing the raw count of each sgRNA by the total count within each of the eight NGS samples (four Cre- and four Cre+). This scaling step adjusts for variations in sequencing depth across samples, allowing for a proportional representation of each sgRNA. To compare sgRNA representation between conditions, the ratio of normalized counts (Cre+/Cre-) was calculated within each of the four biological replicates, providing a quantitative measure of relative sgRNA enrichment or depletion in Cre+ samples. For sgRNA-level analysis, a two-tailed t-test with false discovery rate (FDR) correction was performed on these ratio scores across biological replicates to determine statistical significance. Significantly enriched or depleted sgRNAs were identified based on an adjusted p-value threshold of 0.05. For gene-level analysis, MAGeCK^83^ was used to assess the aggregate effect of multiple sgRNAs targeting the same gene. Raw read counts were used as input, with Cre+ recovered samples designated as the treated group and Cre-samples as the non-treated control.

MAGeCK analysis was performed independently for conditions with and without AAV-α-Syn administration, allowing for the identification of gene-level effects specific to each experimental condition. Each condition included four biological replicates, derived from the pooling of five animals per replicate, as described previously.

### In Vivo CRISPR Validation of Stxbp1

To validate sgRNA-mediated modulation of α-synuclein toxicity, we performed in vivo loss-of-function experiments in Dat-Cre;LSL-Cas9 mice. Mice at two months of age were injected via retro-orbital route with AAV-PHP.eB vectors expressing a U6-driven sgRNA and a Cre-dependent hSyn-DIO-smFP.HA reporter. For the experimental group, the sgRNA targeted Stxbp1, while a non-targeting sgRNA was used in the control group. Each animal received a single sgRNA construct (Stxbp1 or control). Four weeks later, AAV1/2 carrying the human A53T α-synuclein transgene was unilaterally injected into the midbrain to induce neurodegeneration, while the contralateral side served as an internal control.

Three weeks after α-synuclein injection, mice were perfused and brains were coronally sectioned for histological analysis. All immunostaining and subsequent image acquisition were performed in a blinded manner; the experimenter was unaware of mouse genotype or treatment condition throughout the staining and imaging process. Sections were immunostained for tyrosine hydroxylase (TH), and dopaminergic neuron survival was assessed by comparing the number of TH-positive cells between the injected (ipsilateral) and uninjected (contralateral) sides within the same slice. Quantification of TH-positive cells was performed using an unbiased, automated pipeline in Ilastik (v1.4.1rc2), ensuring consistent and objective cell counts across all samples.

### Immunostaining of mouse tissue

Mice were anesthetized with isoflurane and transcardially perfused with PBS containing heparin, followed by 4% paraformaldehyde (PFA). Brains were carefully extracted, post-fixed in 4% PFA with gentle agitation at 4 °C, and sectioned coronally at 40 µm using a vibratome, with hemispheric orientation maintained. Free-floating sections encompassing the substantia nigra pars compacta (SNc) and ventral tegmental area (VTA) were permeabilized, blocked with buffer supplemented with normal goat serum, and incubated overnight at 4 °C with primary antibodies. After washing, sections were incubated with Alexa Fluor–conjugated secondary antibodies at room temperature, protected from light. Stained sections were mounted on slides in the correct right–left order using mounting medium, coverslipped, and stored flat in the dark until imaging.

### Statement on Large Language Model use

ChatGPT (OpenAI) was used solely for grammar and spelling correction during manuscript preparation.

